# Deciphering the scopolamine challenge rat model by preclinical functional MRI

**DOI:** 10.1101/2020.04.08.031534

**Authors:** Gergely Somogyi, Dávid Hlatky, Tamás Spisák, Zsófia Spisák, Gabriella Nyitrai, András Czurkó

**Author notes:** These authors contributed equally to this work (Gergely Somogyi, Dávid Hlatky). <u>Corresponding author</u>: **András Czurkó**, Pharmacological and Drug Safety Research, Gedeon Richter Plc., POB: 27, Budapest 10, H-1475, Hungary.

## Abstract

During preclinical drug testing, the systemic administration of scopolamine (SCO), a cholinergic antagonist, is widely used. However, it suffers important limitations, like non-specific behavioural effects partly due to its peripheral side-effects. Therefore, neuroimaging measures would enhance its translational value. To this end, in Wistar rats, we measured whisker-stimulation induced functional MRI activation after SCO, peripherally acting butylscopolamine (BSCO), or saline administration in a cross-over design. Besides the commonly used gradient-echo echo-planar imaging (GE EPI), we also used an arterial spin labeling method in isoflurane anesthesia. With the GE EPI measurement, SCO decreased the evoked BOLD response in the barrel cortex (BC), while BSCO increased it in the anterior cingulate cortex. In a second experiment, we used GE EPI and spin-echo (SE) EPI sequences in a combined (isoflurane + i.p. dexmedetomidine) anesthesia to account for anesthesia-effects. Here, we also examined the effect of donepezil. In the combined anesthesia, with the GE EPI, SCO decreased the activation in the BC and the inferior colliculus (IC). BSCO reduced the response merely in the IC. Our results revealed that SCO attenuated the evoked BOLD activation in the BC as a probable central effect in both experiments. The likely peripheral vascular actions of SCO with the given fMRI sequences depended on the type of anesthesia or its dose.

## Introduction

On the bases of the cholinergic hypothesis of geriatric memory dysfunction ^1^ scopolamine became a standard reference drug for inducing experimental cognitive impairment in both animals and humans ^2–4^. The so-called “scopolamine challenge” is widely used in preclinical testing of new substances for putative procognitive compounds ^5–7^. This approach, i.e. the reversal of scopolamine-induced impairments, however, has limited predictive validity as it tends to yield a high number of false positives ^8,9^. Specifically, one of the important limitations of the scopolamine (SCO) model is its peripheral side-effects that can influence the learning and memory deficits observed at higher doses of SCO ^4^. In fact, part of the early controversy in using the SCO model was based on the apparent “amnestic” actions of SCO methybromide - a quaternary amine analog of SCO that does not readily enter the brain ^10–13^.

Just the opposite way, by emphasizing the peripheral vascular action of SCO as a primary mechanism, a SCO provocation-based pharmaco-MRI model was proposed as a test for procognitive agents ^5,14^. In this case, another non-blood-brain barrier-penetrating anticholinergic drug, butylscopolamine (BSCO) was used. Indeed, in a pharmaco-MRI study, negative blood-oxygen-level-dependent (BOLD) changes in the prefrontal cortex in isoflurane anesthesia were induced by the i.v. injection of both SCO and BSCO ^14^. However, in α-chloralose anesthesia in a similar preclinical pharmaco-MRI study, increased BOLD contrast in the frontal (orbital) cortex and olfactory nuclei were found after the i.v. injection of SCO while it was absent after BSCO ^15^. These diverging results highlight the difficulty of interpreting preclinical pharmaco-MRI data ^16^ complicated by the use of different anesthetic regimes ^17,18^ that can introduce potential confounds ^19^.

As MRI acquisition parameters could influence the different signal components of BOLD ^20^, contrasting gradient-echo (GE), spin-echo (SE) echo-planar imaging (EPI) and functional arterial spin labeling (ASL) could address interpretation issues and provide a deeper insight into the SCO model. Likewise, performing functional MRI measurements in a block-design in different anesthetic regimes could yield useful, translatable information for the SCO challenge in preclinical drug testing efforts. Therefore, in the present study, we measured whisker stimulation-induced BOLD activation and the effect of SCO in two experiments on an ultrahigh field (9.4T) preclinical MRI system in rats. In the first experiment, SCO and peripherally acting analogue BSCO were used as a pretreatment and BOLD activation were measured to air-puff stimulation of the whisker pad with GE echo-planar imaging (EPI) sequence and with functional ASL in isoflurane anesthesia.

In the second experiment, SCO and BSCO were used in the same way in combined i.p. medetomidine + isoflurane anesthesia and additionally the effect of cholinesterase inhibitor donepezil (DON) was examined on the whisker stimulation evoked BOLD response with both GE EPI and SE EPI sequences.

## Results

For weights, average in-scanner motions, isoflurane doses and respiration rates see Supplementary Materials and Methods (Table S1, S2). These data show that neither of these variables had any effect related to the applied treatment in either experiment.

Our MRI measurements were in a pseudo-randomised cross-over design; therefore, each animal had been measured several times, with a minimum of one-week washout interval between pretreatments and measurements. This way, the number of animals in the given experiment was reduced, and still, at least 15 animals per pretreatment group were measured.

After saline pretreatment, the whisker stimulation evoked BOLD activation was variable in size and depended on the anesthesia and MRI sequences but the barrel cortex (BC) was always involved. An overview of the experiments is shown in Fig. 1 and Table S3.

**Fig. 1.**
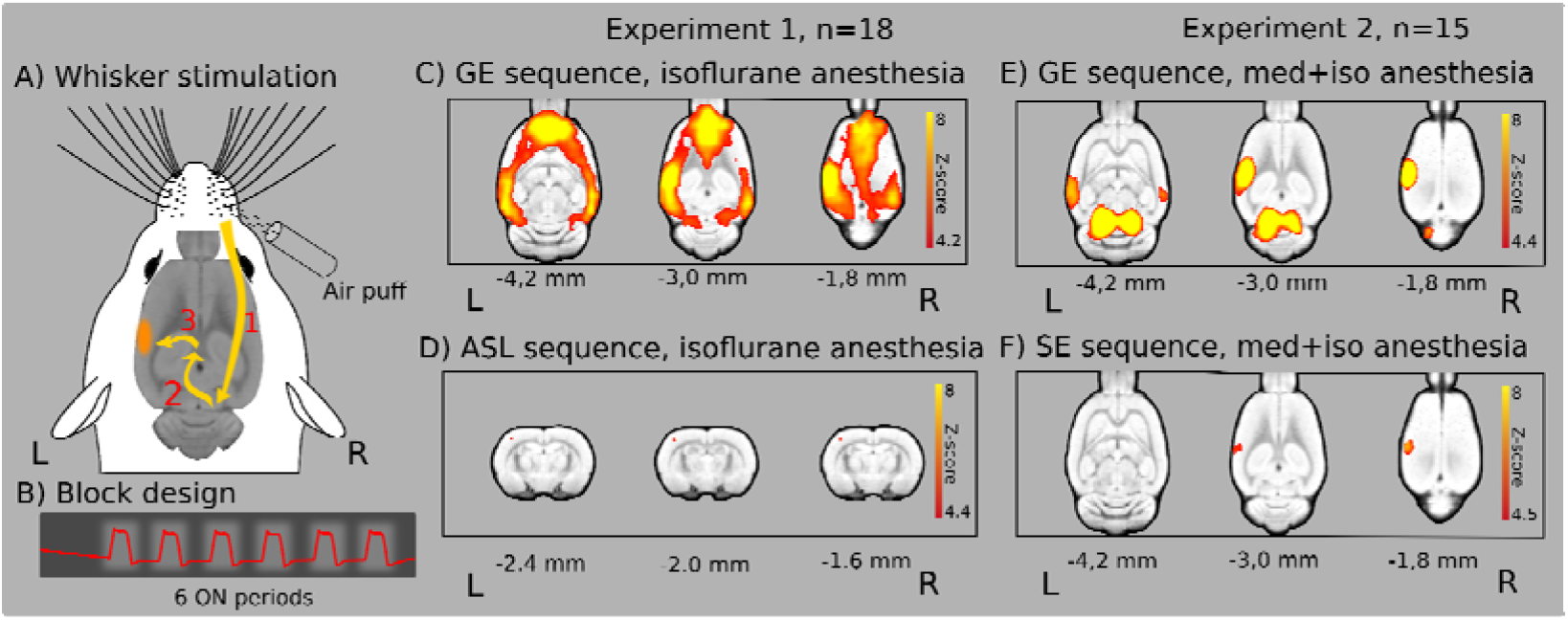
Air-puff stimulation of the right whisker pad in experiment 1 and 2 after saline pretreatment (n=18 and n=15 rats, respectively). *(A)* Signal path from whisker stimulation to activation in the barrel cortex ((1) trigeminal nerve, brain stem, (2) sensory information to the thalamus, (3) the primary somatosensory barrel cortex (BC)). *(B)* Block-design of whisker air-puff stimulation. *(C)* Widespread cortical BOLD activation in isoflurane anesthesia with gradient-echo (GE) EPI sequence. *(D)* Limited activation detected in the BC with the ASL sequence in the same experiment. *(E)* Pronounced activation in the left BC and bilaterally in the IC in the combined anesthesia. *(F)* Limited activation in the BC with the SE sequence in the same experiment. The Z-scores represent voxelwise statistical inference (p<0.05 correcting for multiple comparisons via GRF theory based FWER). The corresponding colorbars are depicted on the right. The z (or y) coordinates (distance from Bregma) are given below the slices (L and R stand for directions).

### Whisker stimulation evoked BOLD responses in isoflurane anesthesia with gradient-echo (GE) EPI sequence

In experiment 1, air-puff stimulation of the right whisker pad evoked widespread cortical BOLD activation in all three treatment groups (Fig. S1 and Table S3). In the saline (Sal) group (Fig. 1c, Fig. 2a), the strongest activation peak was found in the left primary somatosensory cortex, barrel field (L S1 BF). The activation extended bilaterally to wide cortical areas, rostrally it involved the orbital, infralimbic, prelimbic and cingulate cortices, and caudally the auditory, posterior parietal, visual, perirhinal and retrosplenial cortices. For the SCO and BSCO group activations see Fig. S1 and Table S3. For the analysis with three different statistical inferences (voxel, cluster, pTFCE) see Table S4. For local maxima and signal changes with voxelwise statistical inference see Table S5.

**Fig. 2.**
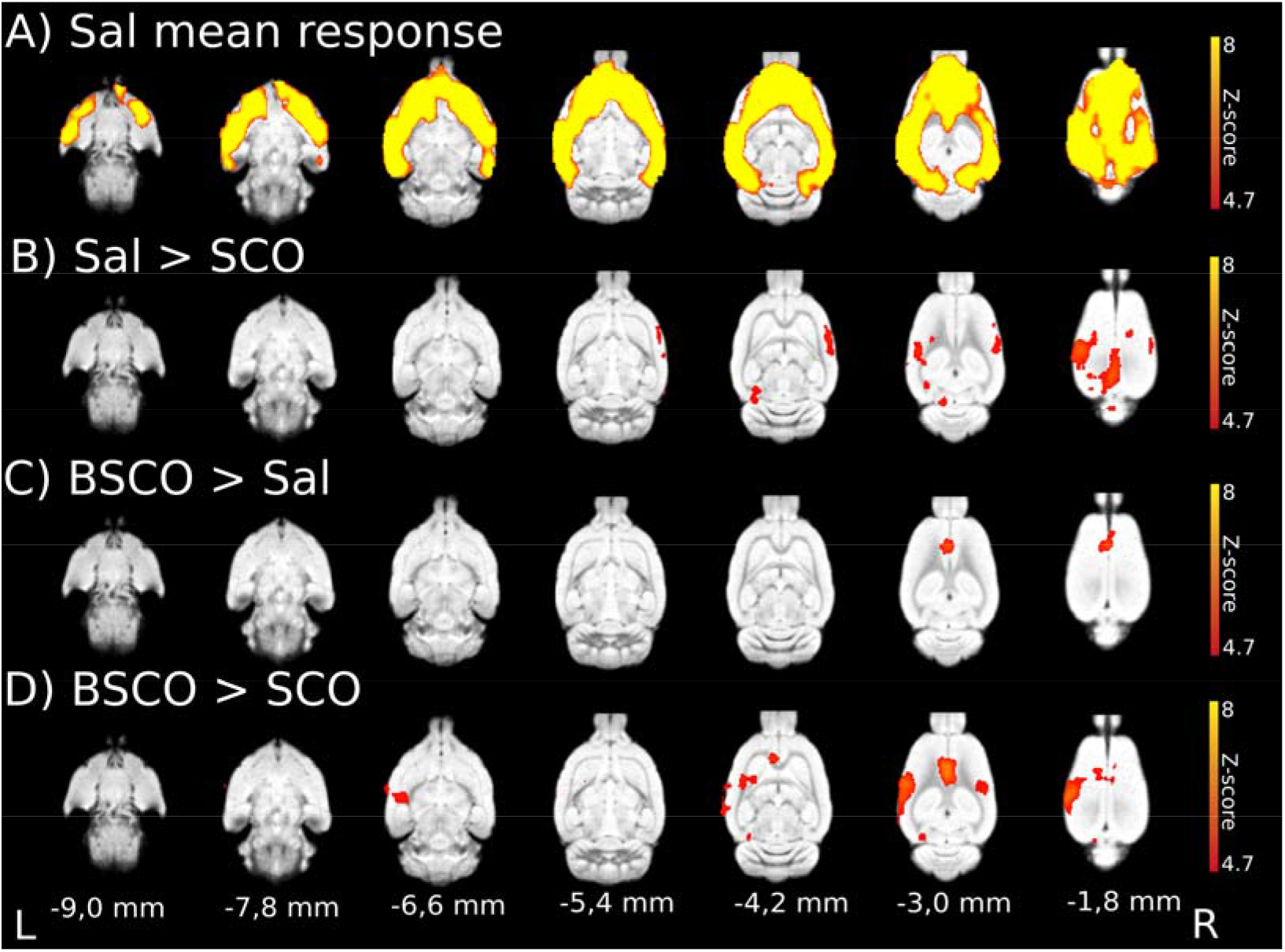
Whisker stimulation evoked BOLD response and evoked BOLD response differences between pretreatment groups with GE EPI sequence in isoflurane anesthesia (n=54; 18 per pretreatment group). *(A)* Mean activation response evoked by whisker sensory stimulation in the saline group. *(B)* Decreased BOLD response in the scopolamine (SCO) treated group compared to saline (Sal > SCO). *(C)* Increased BOLD response in the butylscopolamine (BSCO) group compared to saline (BSCO > Sal). (D) Differences between the SCO and BSCO treated groups (BSCO > SCO). The Z-scores represent probabilistic TFCE (pTFCE) statistics (p<0.05 correcting for multiple comparisons via GRF theory based FWER, red-to-yellow z=4.7 to z=8) from the same data. The corresponding colorbars are depicted on the right. The z coordinates (distance from Bregma) of the selected slices (−9.0 mm; −1.8 mm) are given at the bottom of the figure (L, R stands for directions).

### Treatment group differences in isoflurane anesthesia with GE EPI sequence

Between-group fMRI contrasts revealed statistically significant differences in the groupwise activation patterns between the treatment groups (Fig. 2, Table S6). In the Sal > SCO contrast with probabilistic TFCE (pTFCE; GRF corrected (4.7 < z)) statistics, four areas of statistically significant decreased BOLD response were detected. A large 56.6 mm^3^ cluster in the left primary visual cortex, monocular area (L V1 M) which also contained the L S1 BF (Fig. 2b, Table S6). Besides the L V1 M, the three areas were the secondary somatosensory cortex (R S2; 8.7 mm^3^), the presubiculum (L PrS; 4.1 mm^3^) and the primary auditory cortex (R Au1; 0.4 mm^3^; Fig. 2b, Table S6). With clusterwise statistics, the L S1 BF cluster separated from the L V1 M cluster, and an additional larger cluster in the right primary somatosensory cortex, forelimb region (R S1 FL) was detected (Table S6). For voxelwise statistics see Table S6.

In the SCO > Sal contrast only the uncorrected pTFCE (3.1 < z) statistics revealed a difference in the cingulate cortex (L Cg1, Table S6). In the BSCO > Sal contrast all statistics (Fig. 2c, Table S6) revealed significant difference in the cingulate cortex (L Cg2 or Cg1, Table S6).

In the BSCO > SCO contrast the combination of the Sal > SCO and BSCO > Sal contrasts appeared. All statistics revealed differences in the L S1 BF and L Cg2 (Fig. 2d, Table S6).

There were no significant differences in the Sal > BSCO and SCO > BSCO contrasts except some sub-mm3 in the voxelwise and pTFCE statistics, see Supplementary Table S6 (Table S6).

### Whisker stimulation evoked responses with functional ASL sequence in isoflurane anesthesia

Whisker stimulation-induced activations were focal in the perfusion fMRI analysis of the ASL measurement. In the L S1 BC significant clusters appeared in the Sal (see Fig. 1d) and BSCO treatment groups with all statistical inference (Tables S7), but in the SCO treatment group this focal activation could only be detected with the uncorrected pTFCE (3.1 < z; Fig. S3 and Tables S7) statistic. To define an unbiased somatosensory region-of-interest (ROI), the pooled-group mean BOLD activation (combined Sal, SCO, and BSCO groups) was calculated with clusterwise statistical inference and the average BOLD responses in the ROIs by treatment groups were depicted in boxplots (Fig. S2). Between-group perfusion fMRI analysis of the groupwise activation patterns revealed no statistically significant differences between the treatment groups. For signal changes during functional ASL measurements and the difference between the tagged and non-tagged ASL images over time see Supplementary Fig. S4 and Fig. S5.

### Whisker stimulation evoked BOLD responses in medetomidine-isoflurane combined anesthesia with the GE EPI sequence

In experiment 2, the whisker stimulation-induced BOLD activations had a characteristic focal appearance (Fig. 1e) in all four treatment groups (Fig. S6 and Tables S8). In the Sal group, with the pTFCE (Fig. 3a) as well as with the voxelwise statistics, the strongest activation peaked in the left external cortex of inferior colliculus (L ECIC) but the large activation area bilaterally involved the inferior colliculus (IC) and extended rostrally, bilaterally to the medial geniculate nucleus and on the left, to the ventral posteromedial thalamic nucleus (VPM). A second large activation area peaked in the L S1 BF that also extended to the left auditory cortex and temporal association cortex. The third smaller activation area peaked in the right primary auditory cortex (R Au1).

**Fig. 3.**
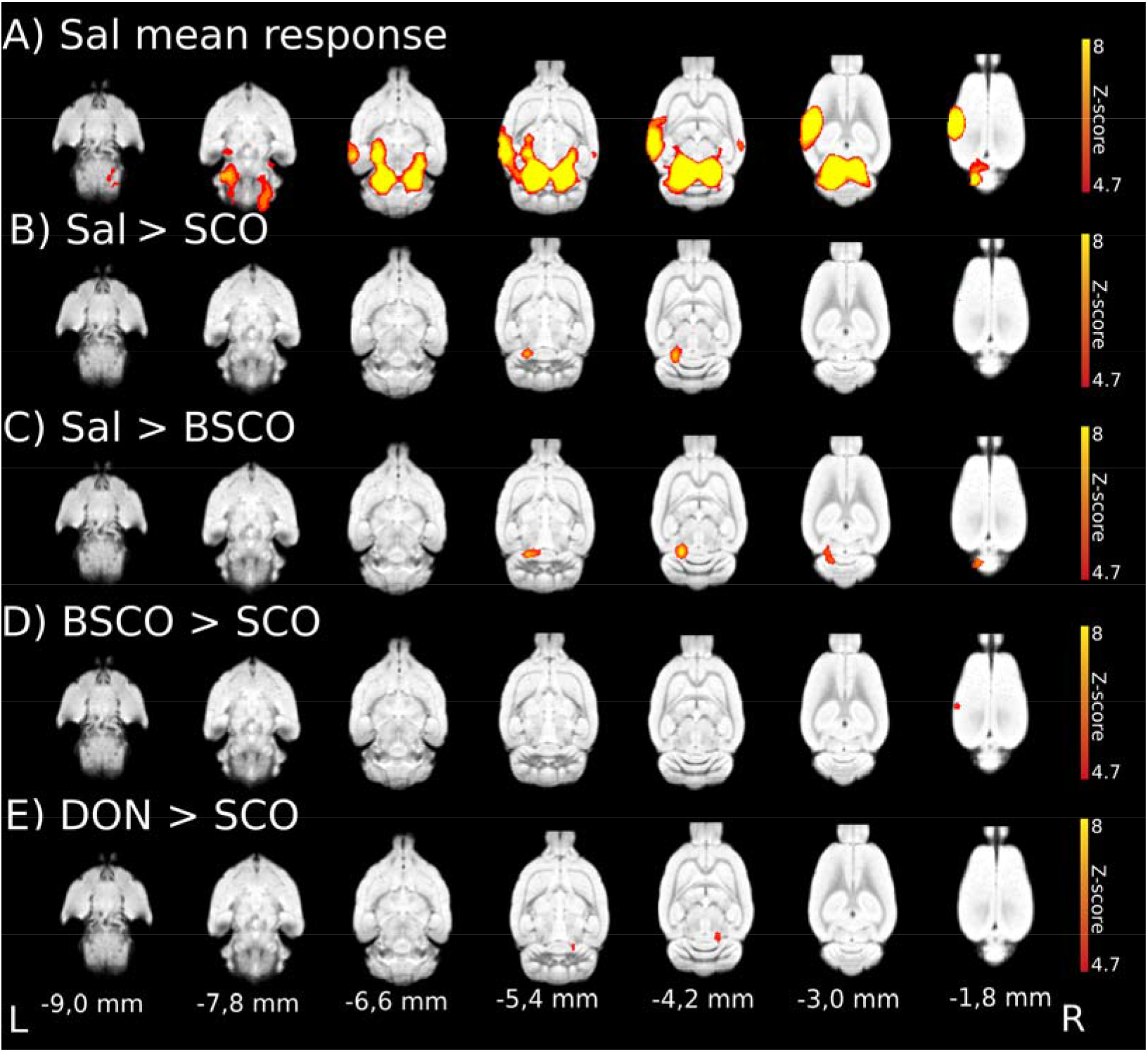
Whisker stimulation evoked BOLD response and evoked BOLD response differences between treatment groups with GE EPI sequence in medetomidine-isoflurane combined anesthesia (n=62; 15, 16, 16, 15 of Sal, SCO, BSCO, DON, respectively). *(A)* Mean activation response evoked by whisker sensory stimulation in the saline (Sal) group. *(B)* Sal > SCO contrast. *(C)* Sal > BSCO contrast. *(D)* BSCO > SCO contrast. *(E)* DON > SCO contrast. Decreased BOLD responses in the BC due to SCO *(B, D)* and in the IC due to both SCO and BSCO *(B,C)*. Increased BOLD responses in the right IC in the DON treatment group compared to the SCO group *(E)*. The Z-scores represent probabilistic TFCE (pTFCE) statistics (p<0.05 correcting for multiple comparisons via GRF theory based FWER, red-to-yellow z=4.7 to z=8) from the same data. The corresponding colorbars are depicted on the right. The z coordinates (distance from Bregma) of the selected slices (−9.0 mm; −1.8 mm) are given at the bottom of the figure (L, R stands for directions).

In the SCO and BSCO groups (1.0 mg/kg) the whisker stimulation-induced activation patterns were very similar to the Sal group (Fig. S6 and Tables S8). In the DON group, the same activation pattern was seen but the activations were somewhat smaller (Fig. S6 and Tables S8).

### Treatment group differences in medetomidine-isoflurane combined anesthesia with the GE EPI sequence

Between-group fMRI analysis contrasts revealed significant differences in this measurement (Fig. 3, Table 1). In the Sal > SCO contrast the pTFCE statistics (GRF corrected (4.7 < z)), revealed decreased activation due to SCO in the L ECIC and one voxel in the L S1 BF (Fig. 3b) while the uncorrected pTFCE (3.1 < z) statistics detected a larger cluster in the L S1 BF and the left and right ECIC (L & R ECIC, Table 1). For voxelwise and clusterwise statistics, see Supplementary Table S10 (Table S10).

**Table 1.**
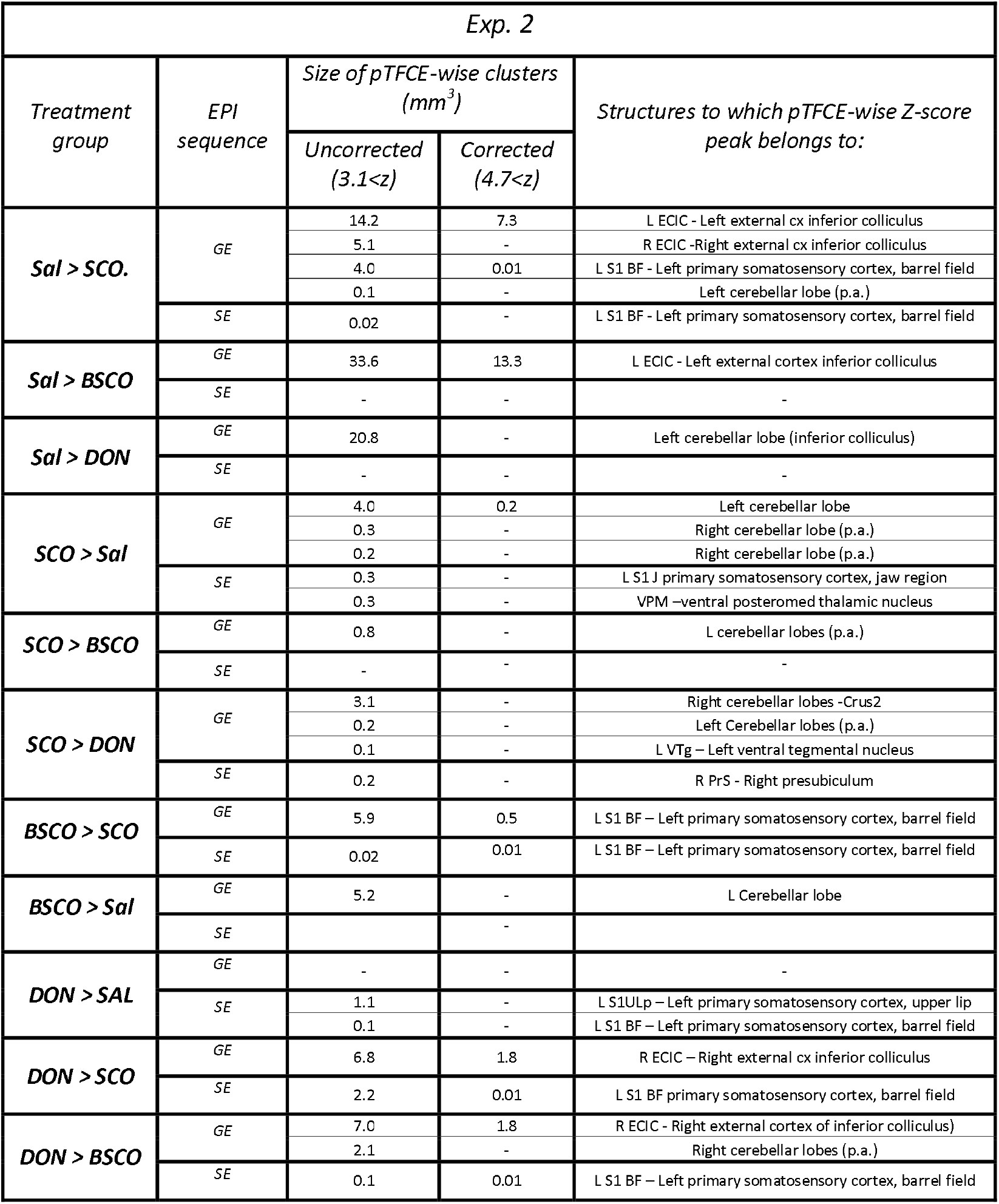
Evoked activation differences in medetomidine-isoflurane combined anesthesia with GE EPI and SE EPI sequences (the size of clusters in mm^3^ and the brain structures to which Z-score (pTFCE statistical inference, p=0.05 via GRF theory base FWER) peaks belong to). N=62; 15, 16, 16, 15 of Sal, SCO, BSCO, DON, respectively. p. a. - stands for probable artefact, as our gradient echo images are not accurate enough around the cerebellum to firmly conclude the difference.

In the Sal > BSCO contrast, pTFCE (GRF corrected (4.7 < z)) statistics revealed a statistically significant decreased BOLD response due to BSCO in the L ECIC (Fig. 3c, Table 1 and Table S10).

In the BSCO > SCO contrast all statistics revealed a difference in the L S1 BF (Fig. 3d, Table 1 and Table S10). Likewise comparing the Sal > SCO and Sal > BSCO contrasts, it might suggest that in this measurement changes in the L ECIC could be a peripheral effect (Fig. 3, Fig. 4, and Table 1).

**Fig. 4.**
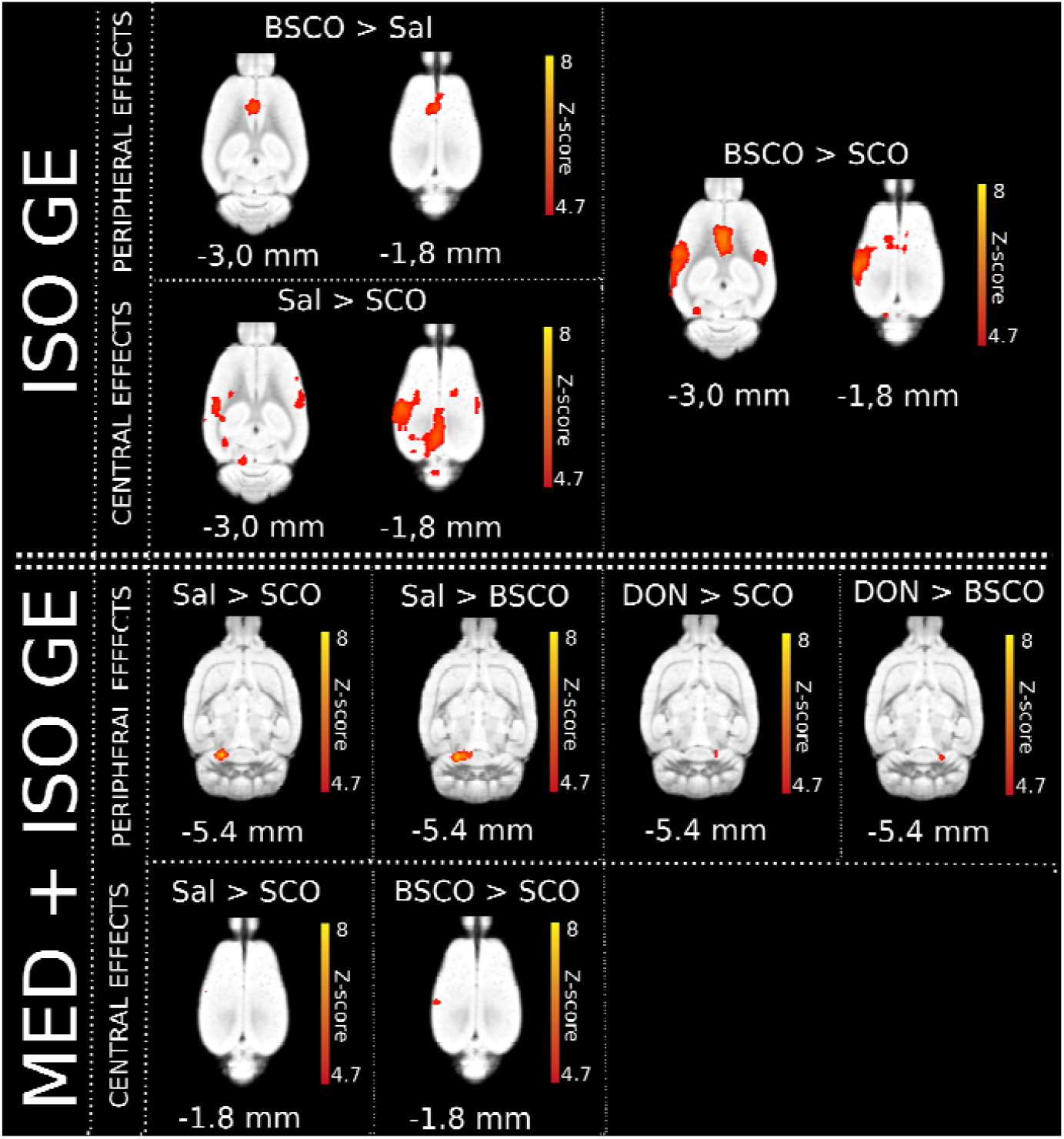
Summary figure of the whisker-stimulated evoked BOLD response differences between pretreatment groups with GE EPI sequences in both anesthesia types (isoflurane – top: ISO GE, (n=54; 18 per pretreatment group); medetomidine-isoflurane – bottom: MED + ISO GE, n=62; 15, 16, 16, 15 of Sal, SCO, BSCO, DON, respectively) according to their peripheral/central effects. In ISO anesthesia, in the BSCO > Sal contrast a peripheral effect, in the Sal > SCO contrast probable central effects are seen. In the BSCO > SCO contrast both effects are seen. In the combined anesthesia (MED + ISO GE) we found probable peripheral effects at the (right or left) inferior colliculus / transverse sinuses in the Sal > SCO, Sal > BSCO, DON > SCO and DON > BSCO contrasts. The central effects (in the left barrel field) are shown in the BSCO > SCO and Sal > SCO group differences. The Z-scores represent the probabilistic TFCE (pTFCE) statistics (p<0.05 correcting for multiple comparisons via GRF theory based FWER, red-to-yellow z=4.7 to z=8). The corresponding colorbars are depicted next to every group difference on the right. The z coordinates (distance from Bregma) of the selected slices (−5.4 mm; −3.0 mm; −1.8 mm) are given at the bottom of the figure.

In both the DON > SCO and DON > BSCO contrasts there were significant differences in the R ECIC area with all statistics (Fig. 3, Fig. 4, Table 1 and Table S10). In the SCO > Sal, SCO > BSCO, SCO > DON and BSCO > Sal contrasts only the uncorrected pTFCE (3.1 < z) statistics found differences (Table 1 and Table S10). In the BSCO > DON and DON > Sal contrasts there were no differences with either statistics (Table S10).

### Whisker stimulation evoked BOLD responses in medetomidine-isoflurane combined anesthesia with SE EPI sequence

In this measurement the whisker stimulation-induced BOLD activations were focal and localized to the L S1 BF in all four treatment groups (Fig. 1f, Fig. S7 and Table S9).

### Treatment group differences in medetomidine-isoflurane combined anesthesia with the SE EPI sequence

Between-group fMRI contrasts revealed significant activation differences only with pTFCE statistics in this measurement, (Table 1). In the BSCO > SCO contrast, with the pTFCE (GRF corrected (4.7 < z)) statistics only one voxel difference was found in the L S1 BF. This one voxel difference also mirrored in the L S1 BF in the DON > SCO and DON > BSCO contrasts (Table 1 and Table S10).

## Discussion

Acute administration of SCO, in several human fMRI studies, induced decreased regional CBF or attenuation in the memory-task-induced increases of regional CBF and BOLD activation in various frontal brain areas using memory and attentional paradigms ^21–25^. These data resemble the SCO-induced hypofrontality in humans ^26^ and rats ^14^ induced by the acute i.v. injection of SCO. The acute i.v. infusion of SCO, however, could transiently produce a reduction of the heart rate and blood pressure ^27^ and this could interfere with the neuroimaging data described above ^28^.

In our neuroimaging study, to avoid the transient effects of SCO, we did not use the traditional preclinical pharmaco-MRI approach with acute i.v. anticholinergic infusion but a “task” fMRI with a pretreatment (1 hour). This pretreatment is similar to those behavioural studies where SCO is used to induce cognitive impairment ^11^. We used the preclinical MRI as a “task” fMRI, by measuring the whisker stimulation-induced BOLD activation in a block design. With this approach, we could not see the dramatic reduction of perfusion or negative BOLD in the prefrontal cortex ^5,14,26^. We also examined the time course of the relative BOLD signal change in the activated barrel field during whisker stimulation. The type of anesthesia and the size of the activated barrel field influenced the relative amplitudes of the BOLD activations (Fig. S10) while the pretreatments not (Fig. S11).

Originally many years before, when we started to investigate the effect of SCO, we surprisingly observed increased BOLD activation in widespread frontal cortical areas due to both SCO and BSCO as a pretreatment (in 1 mg/kg dose; in isoflurane anesthesia) during the whisker stimulation (See Fig. S12, Spisák et al. unpublished data). Similarly, in a previous perfusion imaging study with similar SCO pretreatment (1 hour) an increased rCBV was found in young dogs, unexpectedly ^29^. Furthermore, in the frontal and prefrontal cortical areas during a working memory task hyperactivation due to SCO was also reported ^30^.

One of the limitations of the SCO model is its peripheral side-effects that can influence the learning and memory deficits at higher doses of SCO ^4^. In a preclinical pharmaco-MRI study the 1 mg/kg SCO and BSCO performed similarly in isoflurane anesthesia ^14^, while in α-chloralose anesthesia the 0.8 mg/kg SCO increased the BOLD contrast in the frontal cortex but the same dose of BSCO not ^15^. Importantly, in an eminent translational study of the SCO model, 0.3 mg/kg dose of SCO was used, in rats ^6^. Therefore, in our first experiment, 0.3 mg/kg dose of SCO and BSCO were used, as a pretreatment.

SCO can also increase the acetylcholine release in higher doses ^31,32^ by blocking the M2 autoreceptors ^33,34^ that might as well influence the behavioral and vascular functions. Medetomidine, on the other hand, can suppress the acetylcholine release ^35^ as well as constrict both pial arteries and veins ^36^. Therefore, in our second experiment, in combined medetomidine-isoflurane anesthesia, 1.0 mg/kg dose of SCO and BSCO were used, as a pretreatment.

In experiment 1 (Exp. 1), in isoflurane anesthesia with GE EPI sequence, the whisker stimulation evoked a widespread cortical BOLD activation and group differences revealed a decreased BOLD response in the barrel cortex (BC) due to SCO but not due to BSCO. This SCO result is in line with the human fMRI studies where the SCO typically attenuated the evoked BOLD activation. Interestingly, BSCO increased the evoked BOLD response in the anterior cingulate cortex (Cg1). Only the uncorrected pTFCE statistics detected this effect in case of the SCO pretreatment. This result would mark a peripheral effect on the BOLD activation by BSCO that separated from those of SCO at the 0.3 mg/kg dose, in isoflurane anesthesia.

In Exp. 1, in the ASL-BOLD measurement the whisker stimulation induced small, focal activations in the L BC and the L ECIC (Fig. S2). In the L BC significant clusters appeared in the Sal and BSCO treatment groups, but in SCO this focal activation could only be detected by the uncorrected pTFCE (3.1 < z) inference ^37^. The apparent difference in activation maps between the GE EPI and ASL sequences is probably due to the sequence differences. Our FAIR-ASL sequence used a SE EPI readout with 18 ms echo time with non-negligible T2 weighting, resulting in a mixture of SE-specific BOLD contrast and perfusion contrast. This mixed contrast was utilized during the ASL / Perfusion fMRI image analysis.

The GE EPI sequence is sensitive to both the intra-and extravascular effects of BOLD activation-induced changes while the refocusing pulse in the SE EPI sequence suppresses the extravascular signal around large blood vessels while leaving the signal around the microvasculature intact, thus amplifying its contribution ^38^. The GE BOLD signal, therefore, is more sensitive to large pial surface veins and affected by their local variations ^39,40^.

Exp. 1 was in isoflurane anesthesia that is ideal for non-invasive longitudinal studies ^41–45^. Although isoflurane lacks functional groups and only interacts with its targets through weak polarization forces ^46,47^, there is a chance that it could interfere with the cholinergic system ^48,49^. Isoflurane may also weaken the blood-brain barrier ^50^ by modulating tight junction protein expression ^51^, that may influence the effect of BSCO in our study.

On the other hand, isoflurane anesthesia with spontaneous breathing could dilate the blood vessels ^36,52,53^, and decrease the BOLD responses as well as the spontaneous neural activity, but not the stimulus-evoked neural responses ^53^. This may induce stimulus-evoked propagating slow calcium waves reflected in pan-cortical BOLD fMRI activation ^54^. These may explain the widespread cortical BOLD activation evoked by the whisker stimulation with GE EPI sequence in isoflurane anesthesia in Exp. 1. To address some of the above issues, in Exp. 2 we used combined medetomidine-isoflurane anesthesia, and higher doses of our drug pretreatments as medetomidine can constrict both pial arteries and veins ^36^. Here the same whisker stimulation, with the GE EPI sequence, evoked a characteristic focal BOLD activation. It involved both the left BC and bilaterally the central nuclei of the inferior colliculus (IC). This profound bilateral activation of the IC is an unexpected finding, although the strong background acoustic noise of the scanner is always present. Our fMRI measurement used whisker stimulation in a block design. Therefore, it may look surprising that a constant, background acoustic noise would induce profound activation. It was demonstrated, on the other hand, that the scanner acoustic noise can modulate the somatosensory stimulation. Specifically, the simultaneous acoustic noise facilitated the activation of the somatosensory cortex ^55^. The IC receives ascending projections from both auditory and somatosensory nuclei. Therefore it should be regarded, as a node in multisensory integration ^56,57^. Nonetheless, the profound activation of the IC in the combined medetomidine-isoflurane anesthesia in Exp 2 is still remarkable. Regarding anesthesia, it is also interesting that in an 18F-fluorodeoxyglucose (18F-FDG) PET study, auditory activation resulted in a higher proportion of activated voxels in the IC, but not in the auditory cortex, in combined medetomidine anesthesia than in another type of anesthesia ^58^. Therefore, medetomidine itself might be contributed to the observed profound IC activation.

Group differences with the GE EPI sequence show that both SCO and BSCO decreased the BOLD activation in the IC (Sal > SCO, Sal > BSCO), while SCO also decreased the BOLD activation in the BC. In the BSCO > SCO contrast all statistics revealed the difference in the L S1 BF. These results may suggest that SCO could decrease the BOLD activation in the L S1 BF as through a central effect. Likewise comparing the Sal > SCO and Sal > BSCO contrasts, it might suggest that in this anesthesia the bilaterally decreased activation in the ECIC could be at least partially a peripheral effect that could involve the left and right transverse sinuses. A similar mechanism could explain the decrease in activation in the L ECIC in the Sal > DON contrast, although it localized to the left, activated side. This unilateral effect of DON on the left side can explain the unilateral R ECIC effect in both the DON > SCO and DON > BSCO contrasts.

Therefore, we think that in this anesthesia, we observed both the central (BC) and the peripheral (IC) effects of SCO. With the same GE EPI sequence, we could not reveal a significant difference in the BC, in the DON > SCO contrast. Interestingly, differences in some voxels were detected in the SE EPI measurement by the pTFCE statistical inference.

Our study attempted to preclinically decipher the neuroimaging correlates of the SCO challenge with different statistical inferences. We revealed that the SCO attenuated the evoked somatosensory BOLD activation in the BC, independently from the anesthesia. Its perceived peripheral action depended on the type of anesthesia, dose and fMRI sequences.

The limitation of the study was the lack of the measurement of end-tidal CO2, which plays a pivotal role in vasodilation and the fluctuations of it correlate with fluctuations in global BOLD signal intensity ^59,60^. Additionally, the comparison of SCO challenges under different anaesthesia conditions is limited as different amounts of SCO were used in Exp. 1 & 2.

## Methods

### Animals

Thirty-four drug-naïve adult male Wistar (RJ Han, Harlan Laboratories (Envigo)) rats were used for the two experiments (Exp.1: n=18; Exp.2: n=16,). The animals were kept by controlled laboratory conditions (constant temperature and humidity) in polycarbonate cages under standard in a 12h light/dark cycle with the light on at 6:00 a.m. The rats were fed with conventional laboratory rat food (sniff R/M + H Spezieldiäten GmbH D-59494 Soest), with food and water ad libitum. All the procedures conformed to the guidelines of the National Institutes of Health for the care and use of laboratory animals and were approved by the local Ethical Committee of Gedeon Richter Plc. and were carried out in strict compliance with the European Directive 2010/63/EU regarding the care and use of laboratory animals for experimental procedures. All efforts were made to minimize the number of animals as well as pain or discomfort. Experiments were performed and reported according to the ARRIVE guidelines on animal research.

### Experimental design and drug treatments

The effect of drug pretreatments on the functional MRI activity was examined in a pseudo-randomized cross-over design. On any given measurement day, equal numbers of animals received the various pretreatments of the given experiment. There was a minimum interval of seven days between repeated measurements of the same animal. For the average weight of the animal groups see Supplementary Materials and Methods (Table S1, S2).

In experiment 1 (Exp.1), conscious animals received saline intraperitoneally (i.p.) applied in a volume of 1 ml/kg. or SCO (scopolamine hydrobromide, Tocris) or BSCO (Scopolamine N-butyl bromide, Sigma Aldrich) as pretreatment 1 hour before the start of the MRI measurement. SCO or BSCO were dissolved in saline and injected intraperitoneally at a dose of 0.3 mg/kg (1 ml/kg).

In experiment 2 (Exp.2), conscious animals received saline (1 ml/kg, i.p.), SCO (1 mg/kg), BSCO (1 mg/kg) or DON (Donepezil hydrochloride, 71078800, L77028F, Gedeon Richter Plc.). SCO and BSCO were dissolved in saline and injected in a volume of 1 ml/kg (i.p.). DON was dissolved in distilled water and was injected at a dose of 3 mg/kg, in a volume of 1 ml/kg (i.p.).

### Anesthesia during the fMRI scanning

In Exp.1 after the drug pretreatment animals were anesthetized with 5% isoflurane (Forane, Abbot Laboratories) and then maintained at approximately 1.25% during scanning. Other part of the scanning procedure was identical to the procedures outlined previously^43–45^. In Exp.2 the anesthesia was introduced with a bolus of medetomidine injection (0.05 mg/kg i.p. dexdor, dexmedetomidine, Orion Pharma) plus 5% isoflurane in air and the concentration of isoflurane was lowered to 0.25% - 19 1.5% during scanning with the aim of keeping the respiration rate near 60/min. For mean isoflurane dose of the animal groups see Supplementary Materials and Methods (Table S1, S2).

### Image acquisition

The MRI experiments were performed on a 9.4 T Varian (Agilent) Direct Drive MRI system (Varian Medical Systems Inc., Palo Alto, CA). MRI system parameters described in detail previously^43–45^, also see Supplementary Materials and Methods.

Anatomical images for the fMRI registration were proton density (PD) weighted gradient-echo multi-slice imaging (GEMS) in horizontal orientation, and/or a T2 weighted fast spin-echo multi-slice imaging (FSEMS) anatomical scans in axial orientation, see Supplementary Materials and Methods.

### BOLD fMRI EPI sequences

Echo planar imaging (EPI) sequences were used for the BOLD measurements. Gradient-echo EPI (GE EPI: TR/TE = 1000ms/10ms, FA: 90) and spin-echo EPI (SE EPI: TR/TE = 1000ms/40ms, FA: 90^⍰^) sequences were used (Supplementary Materials and Methods).

### ASL (fMRI) sequence

For multislice arterial spin labeling (ASL) measurements a flow-sensitive alternating inversion recovery (FAIR-ASL, TR/TE = 3.6s/18ms, matrix: 64×64, slice thickness: 2mm, 9 axial slices) sequence was used as described by Nashrallah et al. in^61^, see Supplementary Materials and Methods.

### Whisker stimulation

During the functional MRI experiments, whisker stimulation by air puffs was used in a block-design fashion. The stimulation procedure was identical to the procedure outlined previously^43–45^. In short stimulation was delivered to the right whisker pad of the animals through a tubing system that was integrated into the holding cradle. Air puffs were delivered at a frequency of 1⍰Hz with a blowing time of 200⍰ms. The duration of one stimulation block was 30 s followed by 60 s rest and one measurement session contained 6 blocks in case of the GE-EPI and SE-EPI BOLD fMRI measurements with an initial 120 s resting. For ASL-BOLD fMRI, a 36-72 s on-off cycle was used. See Supplementary Materials and Methods for details.

### Exclusion criteria

Exclusion criteria were fixed before data analysis as follows: scans with striking imaging artifacts (based on visual inspection) or with an average root mean squared relative displacement greater than 0.05⍰mm during the fMRI scan (as calculated by FSL) were subject to exclusion from all analyses.

None of the scans had to be discarded due to artifacts or in-scanner motion issues, resulting in N⍰=⍰54 for the whole-brain fMRI analysis of the first experiment. On the other hand, in the second experiment, 2 measurements had to be excluded because of in-scanner motion and inadequate anesthesia (1 saline and 1 donepezil), resulting in N⍰=⍰62.

## Data analysis

Throughout the image processing procedure, we intended to follow the conventional human-research practice in fMRI analysis. The analysis was carried out in a multistage process using the image analysis software package, FSL^62^ (FMRIB’s Software Library, www.fmrib.ox.ac.uk/fsl) and in-house developed software tools. Small-animal imaging specific details of the applied image processing pipeline were described in detail previously ^43–45^. The present data analysis is detailed in Supplementary Materials and Methods in section Data analysis.

### Functional MRI analysis

Data processing for functional images (GE EPI, SE EPI and ASL –see also Supplementary Materials and Methods) were carried out using FSL FEAT to investigate the effects of treatment groups and their differences. Gaussian kernel (for spatial smoothing and high-pass temporal filtering was applied. After signal prewhitening, the first-level statistical analysis was carried out using the general linear model with local autocorrelation correction ^63^. The analysis-block design convolved with HRF-modelled each rat’s data from each session: one regressor modeling the stimuli (convolved with a double-gamma canonical HRF), its temporal derivate and included 5 noise-ROI-based CompCor ^64^ confounder variables. Individual statistical Z-score images for the BOLD regressor-of-interest were obtained (average BOLD response to the stimuli). Using the spatial transformations computed during image coregistration and standardization, Z-score maps resulting from the individual statistical analysis were realigned to the common standard space to achieve spatial correspondence during the higher-level analysis.

Fixed-effect group-level statistical analysis was performed on the obtained z-stats using FSL FEAT for voxelwise inference using Gaussian random field (GRF) theory based maximum height thresholding with a (corrected) significance threshold of P=0.05 ^65^. This approach is robustly valid, show the mean activation in groups, but often too conservative to find differences between treatment groups because it frequently falling below 5% of family-wise error (FWE) rates ^66^. Similarly, clusterwise thresholding with a corrected significance threshold of P=0.05 at a cluster-forming threshold of p=0.001 (Z>3.1) was performed. This thresholding method was used to enhance the detection of large activation clusters unlikely to be random false positives originating from multiple comparisons ^67^. It gives generally better sensitivity than voxelwise inference for disparity in the evoked responses between groups ^68^. Furthermore, we applied the probabilistic threshold-free cluster enhancement (pTFCE)^37^ with a corrected significance threshold of P=0.05. pTFCE provides a natural adjustment for various signal topologies (thereby enhanced P-values directly, without permutation testing). The method can detect spatially extended but relevant signals of low magnitude, as well as small activations, which remain hidden with the robust, but often too conservative voxelwise statistics. This method was implemented in R software (The R Project for Statistical Computing, https://www.r-project.org).

## Data availability

Data generated and analysed during this study are included in this published article and its Supplementary Information. The raw data datasets generated during the current study are available on https://openneuro.org/datasets/ds003647/versions/1.0.0.

## Acknowledgement

The authors are grateful for the technical assistance of Pálma Diószegi and Katalin Tóthné Fekete. The study was supported by Gedeon Richter Plc., the Hungarian National Research, Development and Innovation Office (KMOP-1.1.5-08-2009-0001), and the National Brain Research Program (KTIA_NAP_13-1-2013-0001; 2017-1.2.1-NKP-2017-00002).

## Author contributions statement

G.S., D.H., T.S., Zs.S., G.Ny. and A.Cz. designed the experiments; D.H. performed the data acquisition; G.S., T.S., Zs.S. analysed the data; G.S., D.H., T.S., G.Ny. and A.Cz. interpreted the data for the work. G.S., D.H. and A.Cz. wrote the main manuscript text and G.S. prepared figures. All authors reviewed the manuscript.

## Additional information

The authors are employees of Gedeon Richter Plc. or were full-time employees of that firm at the time the research was performed, otherwise, they declare no conflicts of interest.

