## Supplementary Material for "Deciphering the scopolamine challenge rat model by preclinical functional MRI"

For the manuscript entitled:

#### Supplementary Materials and Methods

##### Image acquisition

The MRI experiments were performed on a 9.4 T Varian (Agilent) Direct Drive MRI system (Varian Medical Systems Inc., Palo Alto, CA) with a free bore size of 210 mm, containing a 120 mm inner size gradient coil (minimum rise time 140 µs). An actively RF-decoupled quadrature-driven volume coil with an inner diameter of 72 mm (RAPID Biomedical GmbH, Rimpar, Germany) was used for excitation, and a four-channel phased array rat brain coil (RAPID Biomedical GmbH, Rimpar, Germany) located directly above the dorsal surface of the rat's head was used for signal reception. Shimming was done to ensure a water peak linewidth of 18-26 Hz within a 5mm * 5mm * 10mm region inside the brain.

Anatomical images for the fMRI registration were proton density (PD) weighted Gradient Echo multi-slice imaging (GEMS) in the horizontal orientation, and/or a T2 weighted Fast Spin Echo multi-slice imaging (FSEMS) anatomical scans in axial orientation. (GEMS: TR / TE = 500 ms /

4.9 ms, flip angle (FA): 45°, bandwidth: 50 kHz, matrix: 256 * 256, FOV: 35 mm * 35 mm, 36 slices, slice thickness: 0.5 mm, acquisition time: 2 min 10 sec.; FSEMS: TR: 5 s, ETL: 8, effective TE: 33 ms, bandwidth: 40 kHz, matrix: 128 * 128, FOV: 22mm * 22mm, 30 axial slices, slice thickness: 1 mm, acquisition time: 1 min 20 s.)

#### BOLD fMRI EPI sequences

Echo Planar Imaging (EPI) sequences were used for the BOLD measurements. Gradient Echo - Echo Planar Imaging (GE-EPI: TR / TE = 1000 ms / 10 ms, FA: 90°) sequence and 1-shot Spin Echo - Echo Planar Imaging (SE-EPI: TR/TE = 1000 ms / 40ms, FA: 90°) sequence were used in block-design fMRI. In the case of both sequences FOV: 30 mm * 30 mm, slice thickness: 0.8 mm, 17 horizontal slices in an interleaved order, matrix: 38 * 38, resulting in isotropic spatial resolution of 0.8 mm. 660 images were acquired for a total acquisition time of 11 min. In Experiment 1 and Experiment 2 for both GE and SE sequences even-odd image pairs were acquired with opposite readout gradient direction and were combined for ghost removal (van der Zwaag et al., 2009) resulting in 330 images and an effective time resolution of 2 s for further analysis. While the BOLD signal has optimal contrast-to-noise ratio near an echo time of 18 ms at 9.4T with GE-EPI, a 10 ms echo time reduces signal dropouts due to B0 field inhomogeneity in certain parts of the rat brain. Since the Ernst-angle is about 50-55° in the current settings, using a flip angle of 90° introduces T1 weighting resulting in an added non-BOLD contrast, possibly by enhancing local vascular contributions, notably, the changes in CBF and CBV (Gao & Liu, 2012).

#### ASL (fMRI) sequence

For cerebral blood flow (CBF) measurements multislice arterial spin labeling with flow-sensitive alternating inversion recovery (FAIR-ASL) sequence was used as described by Nashrallah et al. (Nasrallah, Lee, & Chuang, 2012), but the inversion bandwidth was modified to 10 kHz. Sequence summary: hyperbolic secant adiabatic inversion pulse of 20 ms and 10 kHz, followed by inversion time of 1.2 s. Multislice image acquisition was done by a single-shot SE-EPI scheme with TR/TE = 3.6 s / 18 ms, bandwidth: 250 kHz, matrix: 64 * 64, FOV: 22 mm * 22 mm, slice thickness: 2 mm, 9 slices were acquired in an anterior-posterior direction. A total of 50 tag-control pairs were acquired over 6 min 30 s. It should be noted that the use of the

SE-EPI readout with 18 ms echo time at 9.4T results in non-negligible T2 weighting in the rat brain, and thus a BOLD contrast appears alongside the functional CBF changes with this sequence. This BOLD contrast was utilized during the image analysis accordingly.

##### Whisker stimulation

During the functional MRI experiments, the pneumatic "air-puffed" whisker stimulation was used in a block-design fashion. The stimulation was delivered to the right whisker pad of the animals through a tubing system that was integrated into the holding cradle. Air delivery timing was controlled with a magnetic valve operated by an in-house written MATLAB script (version 2009b, The MathWorks, Inc., Natick, Massachusetts, United States) via an NI USB-6215 Multifunction I/O Device (National Instruments, Austin, USA). The air pressure was set to 1 bar. Air puffs were delivered at a frequency of 1 Hz with a blowing time of 200 ms. The duration of one stimulation block was 30 s followed by 60 s rest and one measurement session, contained 6 blocks in case of the GE-EPI and SE-EPI BOLD fMRI measurements with an initial 120 s resting period after the start of imaging (660 s). In the case of the ASL-BOLD fMRI measurement, one block consisted of 36 s stimulation and a 72 s resting period. At the start of imaging, a ~60 s resting period was set. ASL measurements also contained 6 blocks: 3 consecutive blocks in two ASL measurement sessions (2 x 396 s).

#### Data analysis

Throughout the image processing procedure, we intended to follow the conventional human practice in fMRI analysis. Small-animal imaging specific details of the applied image processing pipeline are below described in detail.

##### Preprocessing

The raw images were converted to NIfTI-format (Neuroimaging Informatics Technology Initiative) by an in-house written MATLAB script (version 2009b, The MathWorks, Inc., Natick, Massachusetts, United States). The anatomical and functional images were reoriented to match the standard orientation of the digitalized version of the Rat Brain Atlas of Paxinos and

Watson (Paxinos & Watson, 2005). All images were rescaled by a factor of ten to achieve image dimensions similar to human data and thus facilitate the use of image processing algorithms developed for human image analysis (Spisak et al., 2017). All the approaches performed on the upscaled images were scale-invariant and does not involve any information loss. The analysis was carried out in a multistage process using the image analysis software package, FSL (Jenkinson, Beckmann, Behrens, Woolrich, & Smith, 2012) (FMRIB's Software Library, www.fmrib.ox.ac.uk/fsl), and in-house developed software tools. FSL SUSAN (Stephen M. Smith & Brady, 1997) was used on structural images for noise reduction with nonlinear filtering to reduce noise whilst preserving the underlying structure. All the fMRI (GE, SE, and ASL) time-series were motion-corrected using MCFLIRT (Jenkinson, Bannister, Brady, & Smith, 2002). FSL BET (Brain Extraction Tool) (S. M. Smith, 2002) was used to remove non-brain areas from the structural and functional images. The fractional intensity threshold was set to 0.7 and the vertical gradient in the fractional threshold was set to 0.1 for functional images. For structural images, the fractional intensity was 0.25 or 0.2 and the vertical gradient was 0.6 or 0 (based on the resolution).

To achieve spatial correspondence for the group analysis, all images were spatially standardized in two-steps (two-level registration). The functional (low-resolution) images were registered to high-resolution structural images using FSL FLIRT (Jenkinson & Smith, 2001), which utilized a 6-parameter rigid-body transformation. The high-resolution structural image was fitted to an in-house developed standard template (Spisak et al., 2017) (average of 200 nonlinearly fitted structural images) by an affine transformation (FLIRT) and a nonlinear deformation field estimated by FNIRT (Andersson, Jenkinson, & Smith, 2007). In the latter coregistration procedure, a 10 x 10 x 10 $mm^3$ warping field (in upscaled space) was estimated with three iterations using a relatively conservative lambda parameter set of 60, 40, and 20. This stricter regularization was applied as the morphological diversity of the rat brain is smaller compared to humans.

#### Functional MRI (fMRI) analysis

##### Gradient echo (GE) & Spin echo (SE)

Before the statistical analysis, the functional GE and SE images were spatially smoothed using a Gaussian kernel of 12.0 mm FWHM (note that the images were upscaled). High-pass temporal filtering (Gaussian-weighted least-squares straight line fitting, with sigma=162.0 s) was then applied to remove slow drifts from the signal. The fMRI data processing was carried out using FEAT (FMRI Expert Analysis Tool) Version 6.00, part of FSL. After signal prewhitening (FSL FILM), the first-level statistical analysis on the time-series was carried out using the general linear model with local autocorrelation correction (Woolrich, Ripley, Brady, & Smith, 2001). The analysis modeled each individual rat's data from each session and included 7 explanatory variables in total: one regressor modeling the stimuli (convolved with a double-gamma canonical HRF), its temporal derivate, to account for slight differences in timing and 5 noise-ROI-based, CompCor (Behzadi, Restom, Liau, & Liu, 2007) confounder variables. For each fMRI session, a noise ROI was delineated based on the temporal signal-to-noise ratio of the BOLD signals (the upper 2 percentiles of within-brain voxels were chosen on each slice), and the first five principal components of the corresponding time-series were extracted from the data, following the broadly used t-CompCor technique.

Individual statistical Z-score images for the first regressor were obtained (average BOLD response to the stimuli). Using the spatial transformations computed during image coregistration and standardization, Z-score maps resulting from the individual statistical analysis were realigned to the common standard space to achieve spatial correspondence during the group-level analysis.

Resulting Z-score maps (both in Experiment 1 and Experiment 2) were processed in three ways for investigating the effect of individual treatment groups (saline, scopolamine, butyl-scopolamine, donepezil) and the differences between groups with a general linear model (where the factor-of-interest was the treatment). First, second-level fixed-effect statistical analysis was performed using FSL FEAT for voxelwise inference using GRF-theory-based maximum height thresholding with a (corrected) significance threshold of P=0.05 (Worsley, 2001). This approach is robustly valid, show the mean activation in groups, but often too conservative to find differences between treatment groups because it frequently falling below 5% of family-wise error (FWE) rates (Eklund, Nichols, & Knutsson, 2016). Similarly, for clusterwise thresholding with a corrected significance threshold of P=0.05 and cluster holding threshold of Z>3.1 was used to enhance the detection of large activation clusters unlikely to be random false positives originating from multiple comparisons (Friston, Worsley, Frackowiak, Mazziotta, & Evans, 1994). It gives generally better sensitivity than voxelwise inference for the disparity in the evoked responses between groups (S. M. Smith & Nichols,

2009). Furthermore, we applied the probabilistic threshold-free cluster enhancement (pTFCE) (Spisak et al., 2018) with corrected significance voxel threshold of P=0.05. In order to illustrate our results thoroughly, in some cases, we also show uncorrected pTFCE statistical inference (we thresholded our enhanced Z-score images with uncorrected $p < 0.001$ value, which corresponds to Z>3.1). This method was carried out with R software (The R Project for Statistical Computing, https://www.r-project.org) which provides a natural adjustment for various signal topologies (thereby enhanced P-values directly without permutation testing).

#### Arterial spin labeling (ASL) / Perfusion fMRI analysis

Data processing for the ASL measurement was carried out using FSL FEAT in order to investigate the effects of treatments (saline, scopolamine, butyl-scopolamine) and their differences. It is possible to analyse perfusion fMRI data (often also referred to as ASL) with FEAT. The data needs special treatment because each voxel's timeseries alternates between "tag" and "control" conditions, with control timepoints having higher intensity. Our FAIR-ASL fMRI sequence contained a relatively long echo time (18ms) therefore there was BOLD signal present that could be modelled as the average of the "tag" and "control" conditions with FEAT in FSL. The perfusion (flow) signal can be modelled as the difference between the tag and control conditions; when there is activation, this difference increases. (https://fsl.fmrib.ox.ac.uk/fsl/fslwiki/FEAT/UserGuide#Perfusion_FMRI_Analysis).

A Gaussian kernel of 12.0 mm FWHM (the images were upscaled to the human-sized brain) for spatial smoothing was used. High-pass temporal filtering (Gaussian-weighted least-squares straight-line fitting, with sigma=162 s) was then applied. After signal prewhitening, the first-level statistical analysis was carried out using the general linear model with local autocorrelation correction. We analysed our data with full perfusion signal modelling, where separate explanatory variables (EVs) model the BOLD signal, the (constant-height) tag-control difference and the modulation of this by the activation. The analysis modeled each individual rat's data from each session and included 11 explanatory variables in total: one regressor modeling the control-tag baseline, one for the stimuli (convolved with a double-gamma canonical HRF), and one regressor is formed by multiplying these two; their temporal derivate (3 regressors) to account for slight differences in timing; 5 noise-ROI-based, CompCor (similar to GE, SE) confounder variables.

Statistical Z-score images for the third regressor were obtained (average BOLD response to the stimuli). Using the spatial transformations (computed beforehand) Z-score maps resulting from the individual statistical analysis were realigned to the common standard. The second-level analysis utilizes the Z-score maps from the first-level evaluation and models the dependence of images by rats (due to imaging method 2 images with 3-3 blocks are generated by rats). The resulting COPE's (contrast of parameter estimates) are used in the third-level fixed-effect analysis as input data. Similar to GE, SE analysis voxelwise inference (using GRF-theory-based maximum height thresholding with a corrected significance threshold of P=0.05), clusterwise thresholding (with a corrected significance threshold of P=0.05 and cluster holding threshold of Z>3.1), and pTFCE methods were applied.

#### Signal Change

The signal change is defined as the ratio of the measured signal during stimulation and resting blocks. The average of all intensities during stimulation (GE and SE: 6 * 30 s; ASL: 6* 36 s) divided by the average of all intensities during resting (GE and SE: ~120 s + 6 * 60 s; ASL: 60 s + 6 * 72 s) resulted in the signal change which is expressed in percentage (%). In our study, we show the signal change of voxels and brain areas (anatomical regions or region-of-interests – where the signal change by voxels was averaged) as well.

#### Relative BOLD change

The relative Blood-Oxygenation-Level-Dependent (BOLD) change is defined as the change of the signal compared to the average starting resting period (120 seconds). We have only examined this effect in the left barrel cortex in the isoflurane anesthesia (only in the gradient echo sequence) and in the combined anesthesia (GE, SE sequences as well), to investigate the effect of the whisker stimulation. After averaging the timepoints a whole-brain average and dispersion were calculated. The whole-brain average was subtracted from the voxel and it was divided by the dispersion. The relative BOLD change can show how much the anesthesia and/or the treatments affect the signal.

#### Supplementary Figures

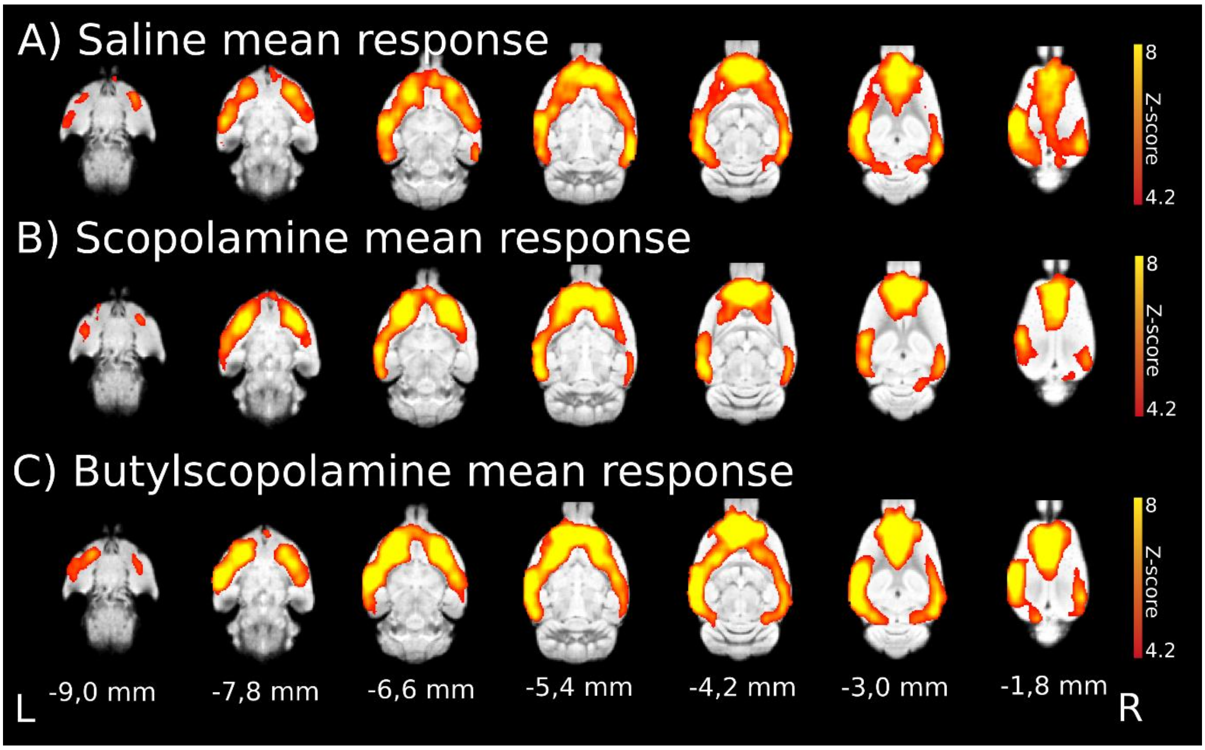


**Suppl. Fig. 1. Whisker stimulation evoked gradient-echo EPI sequence BOLD responses in isoflurane anesthesia and the effect of anticholinergic modulation**. **a** Mean activation response evoked by whisker sensory stimulation in the saline group. **b** Mean activation response in the scopolamine group. **c** Mean activation response in the butylscopolamine group. The z coordinates (distance from bregma) of the selected slices are given at the bottom of the figure (the L and R stand for the left and right directions). The Z-scores represent voxelwise ($p<0.05$, correcting for multiple comparisons via voxel significance) statistical inference (red-to-yellow from z=4.2 to z=8), the corresponding colorbars depicted on the right.

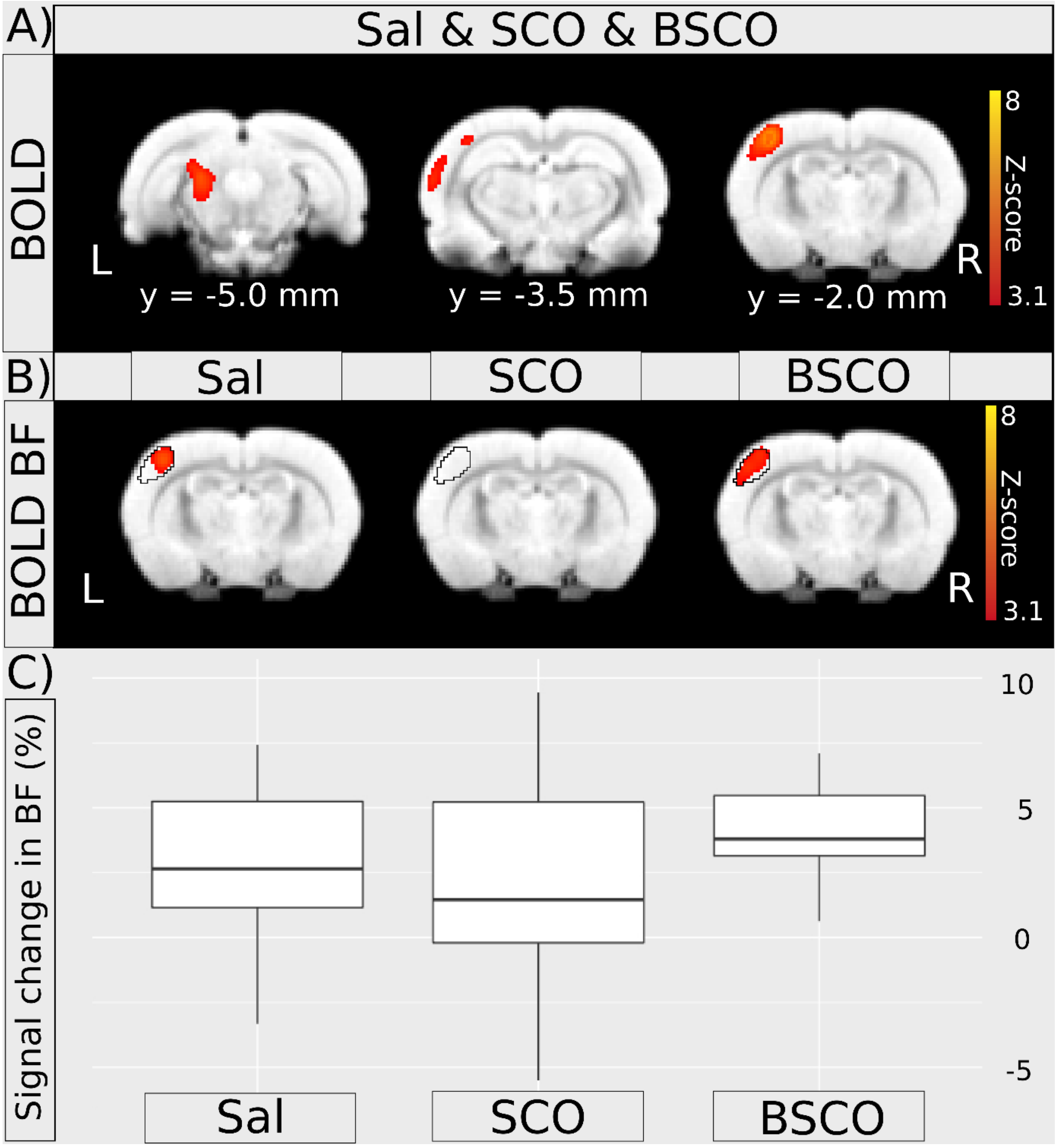


**Suppl. Fig. 2. Whisker stimulation evoked perfusion FMRI analysis of BOLD responses in isoflurane anesthesia with an ASL sequence and the signal changes in the somatosensory barrel cortex (BF). a** Pooled group mean BOLD activation evoked by whisker stimulation (Sal & SCO & BSCO). The y coordinates (distance from bregma) of the selected slices are given at the bottom of the figure (the L and R stand for the left and right directions). **b** BOLD response in the barrel field (BF) by treatment groups and the unbiased somatosensory region-of-interest (ROI) defined from the pooled-group mean activation Z-score > 9 delimited by a black contour. The selected slice y=-2.0; R and L stand for directions. The Z-scores represent clusterwise ($p<0.05$, correcting for multiple comparisons via cluster significance, cluster forming threshold of $z=3.1$) statistical inference (red-to-yellow from $z=3.1$ to $z=8$), the corresponding colorbars depicted on the right. **c** Boxplots represent the average BOLD response in the unbiased somatosensory ROIs (in b) by treatment groups of the 54 ASL measurement. The difference in the means and deviations may explain that there is no significant cluster in the BF the SCO group. The scale of signal change is depicted on the right and is given in percentage (%).

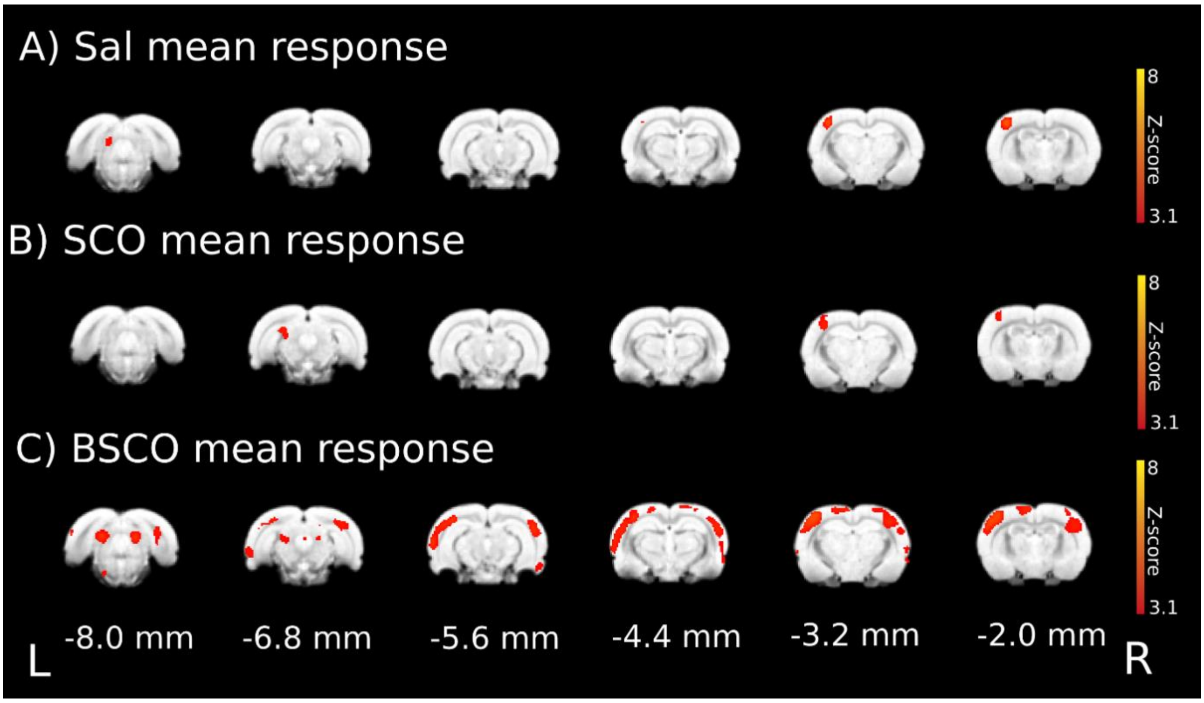


**Suppl. Fig. 3. Whisker stimulation evoked functional responses with an ASL sequence in isoflurane anesthesia (uncorrected pTFCE statistics). a** Mean evoked activation in the control (saline) group. **b** Mean evoked activation in the scopolamine group. **c** Mean evoked activation in the butylscopolamine group. The distance of the coronal slices from bregma (the y coordinates) are given at the bottom of the figure. Z-scores of uncorrected pTFCE statistical inference (enhanced Z-score thresholded at $p < 0.001$ value, which corresponds to $z>3.1$) depicted on the right (red-to-yellow from $z=3.1$ to $z=8$).

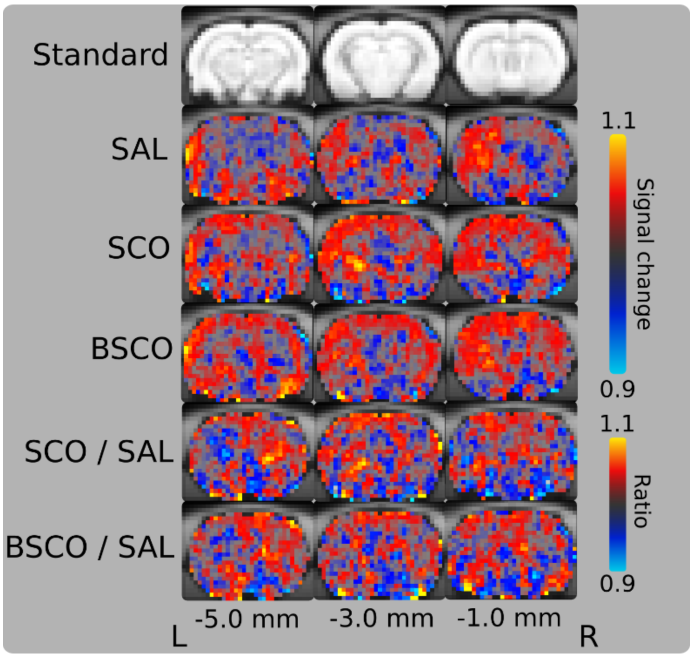


**Suppl. Fig. 4. The signal change during ASL measurement in three slices.** Saline, scopolamine, butylscopolamine, and the two bottom lines illustrate the ratio between scopolamine and saline, butylscopolamine and saline. The z slices distance from bregma are shown in the bottom of the figure (z = -1.0 mm, z = -3.0 mm, z = -5.0 mm; L,R stand for left and right direction). The colorbar shows the signal change or the ratio to control (typically is between 0.9 and 1.1).

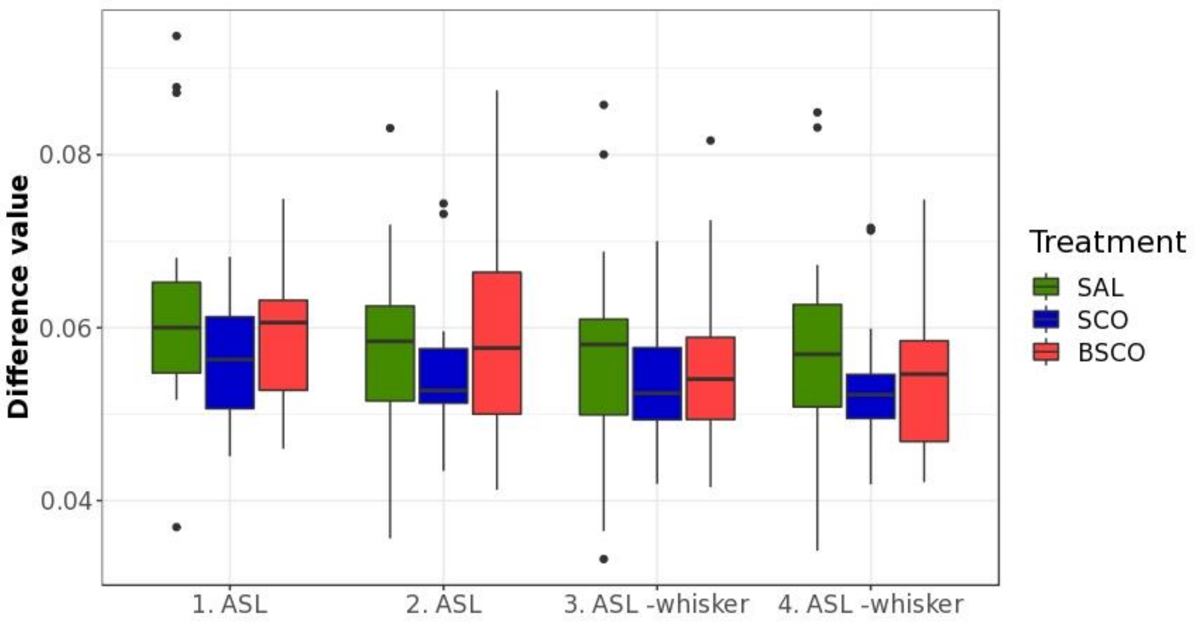


**Suppl. Fig. 5. The difference between tagged and non-tagged images of every ASL measurement (in experiment 1) by treatment groups.** The difference values were obtained from each voxel of ASL images: the non-tagged ones are subtracted from the tagged ones and divided by the base non-tagged. The difference values are averaged per animal and then averaged by every timepoint. The timepoints are the four different ASL measurements (two without and two with whisker stimulation). The green boxplots are the control (SAL) group, the blue ones are the SCO group and the red ones stand for BSCO. Due to this consideration, the influential effect of anesthesia (the effect of isoflurane) -the four time points are within about 25 minutes- can be excluded.

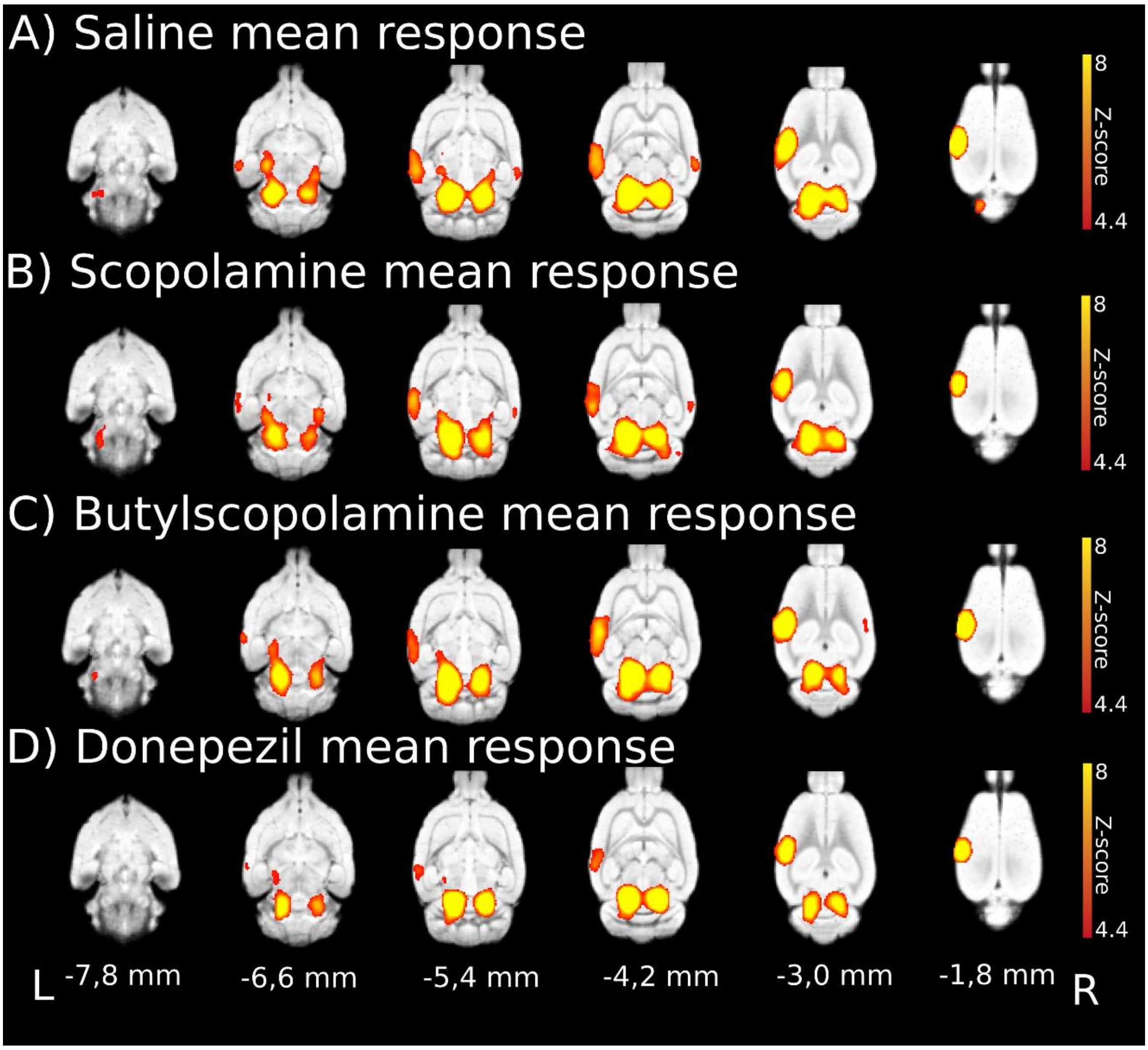


**Suppl. Fig. 6. Whisker stimulation evoked gradient-echo EPI sequence BOLD responses in medetomidine-isoflurane combined anesthesia and the effect of cholinergic modulation. a** Mean activation evoked by whisker sensory stimulation in control (saline) group; **b** mean activation in scopolamine; **c** in butylscopolamine and **d** in donepezil group with gradient-echo EPI sequence. The z coordinates (distance from bregma) and directions (left - L, right - R) are given at the bottom of the figure. Red-to-yellow Z-scores from voxelwise statistical inference ($p<0.05$, correcting for multiple comparisons via voxel significance) are shown on the right.

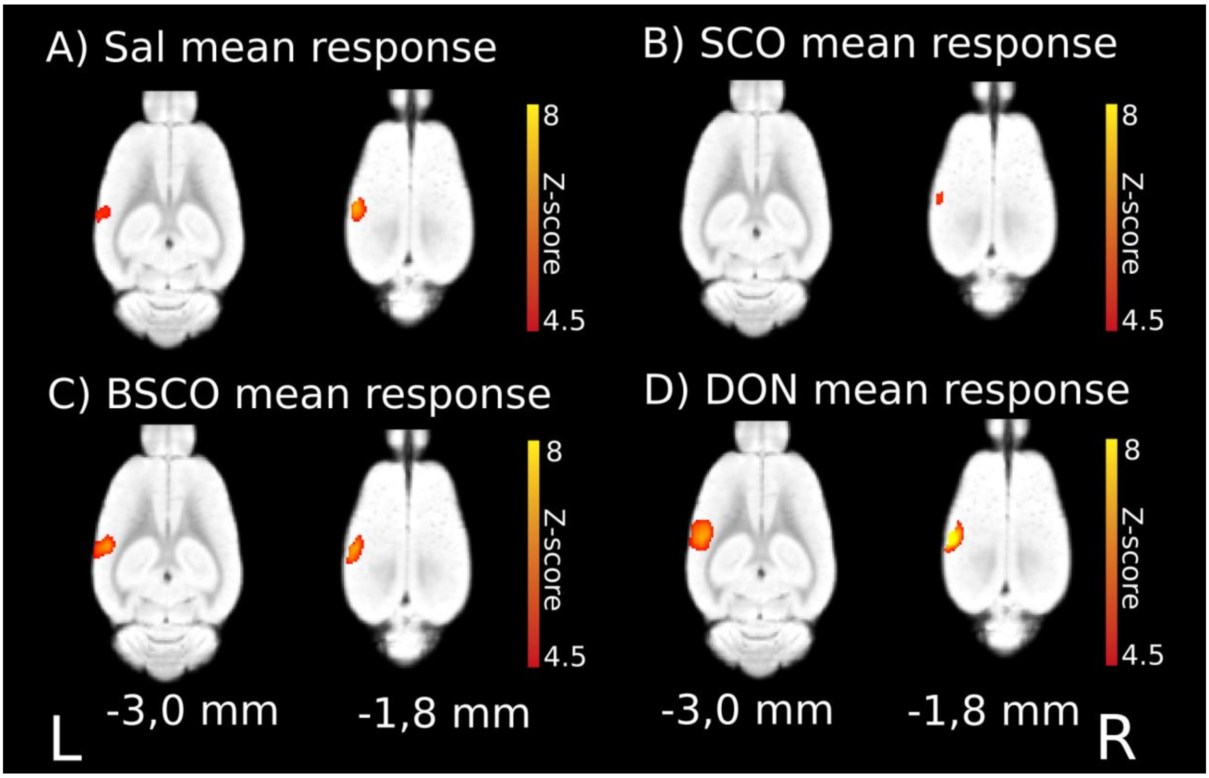


**Suppl. Fig. 7. Whisker stimulation evoked spin-echo EPI sequence BOLD responses in medetomidine-isoflurane combined anesthesia.** Mean activation evoked by whisker sensory stimulation in the barrel field (two slices are shown (distance from bregma): z = -3,0 mm, z = -1,8 mm). **a** saline, **b** scopolamine, **c** butylscopolamine and **d** donepezil group averages. At the bottom of the figure, the L and R stand for the left and right directions. The Z-scores represent voxelwise statistics (red-to-yellow from z=4.5 to z=8, p<0.05, correcting for multiple comparisons via voxel significance), the corresponding colorbar depicted on the right.

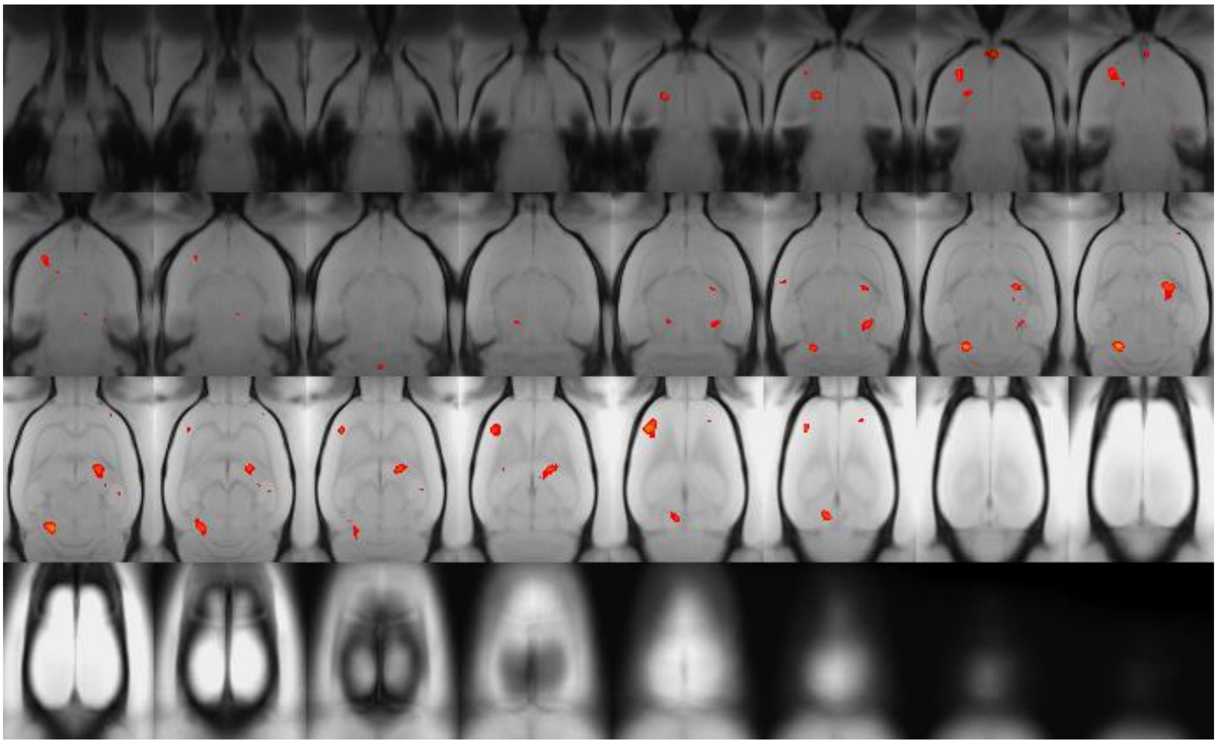

**Suppl. Fig. 8. Whisker stimulation evoked differences between gradient-echo (GE) and spin-echo (SE) EPI sequences in Sal > SCO contrasts of BOLD responses in medetomidine-isoflurane combined anesthesia.** The Z-scores represent cluster statistics ($p<0.05$ correcting for multiple comparisons via GRF theory based FWER and a cluster forming threshold of $p=0.01$, red-to-yellow $z=2.3$ to $z=8$). These results show that there is a difference between the sequence contrast (GE > SE) in various small clusters.

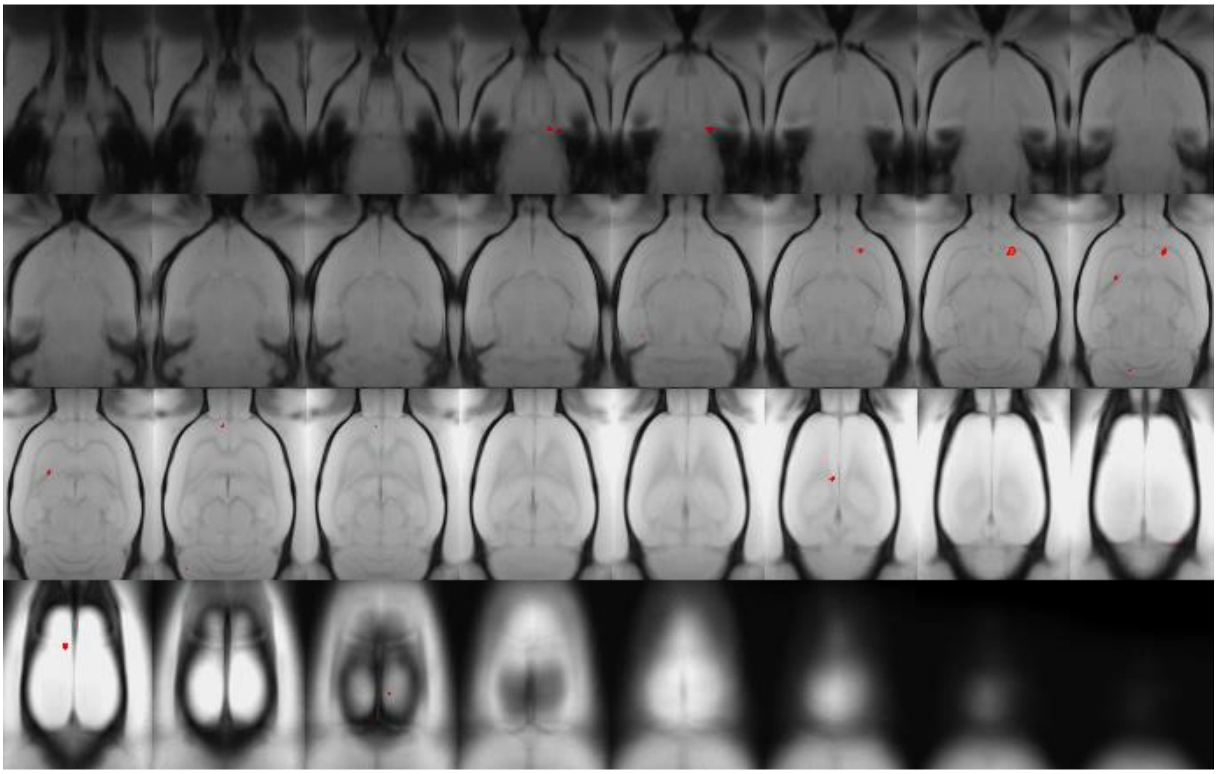

**Suppl. Fig. 8. Whisker stimulation evoked differences between gradient-echo (GE) and spin-echo (SE) EPI sequences in SCO > Sal contrasts of BOLD responses in medetomidine-isoflurane combined anesthesia.** The Z-scores represent cluster statistics ($p<0.05$ correcting for multiple comparisons via GRF theory based FWER and a cluster forming threshold of $p=0.01$, red-to-yellow $z=2.3$ to $z=8$).

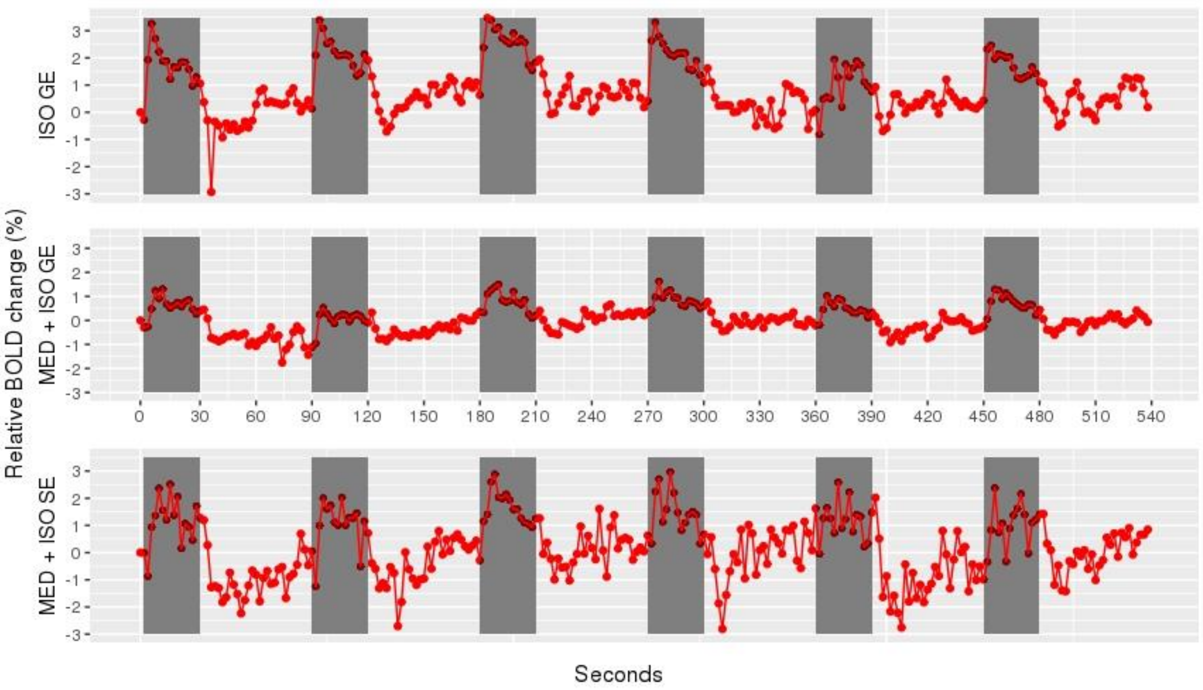


**Suppl. Fig. 10. Time course of relative BOLD activation due to the six whisker stimulation blocks in the average activated barrel field (where the Z-score > 12; ISO GE: 10,38 mm$^3$; MED + ISO: 15,29 mm$^3$; MED + ISO SE 1,56 mm$^3$), for all EPI measurements.** Top row: isoflurane (ISO) anesthesia GE sequence; middle row: combined anesthesia (MED + ISO) GE sequence; bottom row: MED + ISO anesthesia SE sequence. Every timepoint is normalized to the starting point (0. second), which is the average of the signal before the first whisker stimulation (~120 seconds). The y axis shows the relative BOLD change in percentage (%) compared to the initial resting period. The x axis shows the elapsed time in seconds. The 30 seconds long stimulation periods are darkened in the figure.

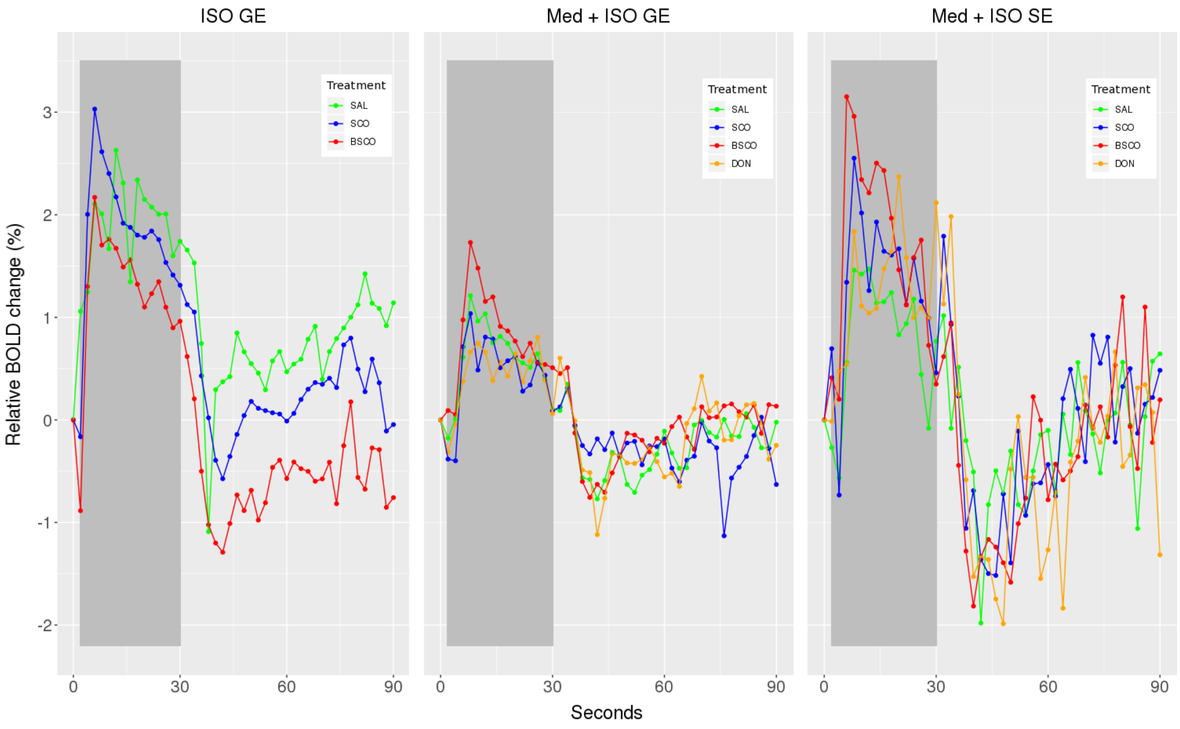


**Suppl. Fig. 11. Relative BOLD change averaged into one block in the average activated barrel field (where the Z-score > 12).** Left figure: isoflurane (ISO) anesthesia GE sequence (area: 10,38 mm$^3$); middle figure: combined anesthesia (MED + ISO) GE sequence (area: 15,29 mm$^3$); right figure: MED + ISO anesthesia SE sequence (area: 1,56 mm$^3$). Every timepoint is normalized to the starting point (0 second the starting point is the average signal of about 120 seconds before the first whisker stimulation) and averaged into one block (30 seconds on and 60 seconds off period). The y axis shows the relative BOLD change in percentage (%) compared to the initial resting period. The x axis shows the elapsed time in seconds. The 30 seconds long stimulation period is darkened in the figure.

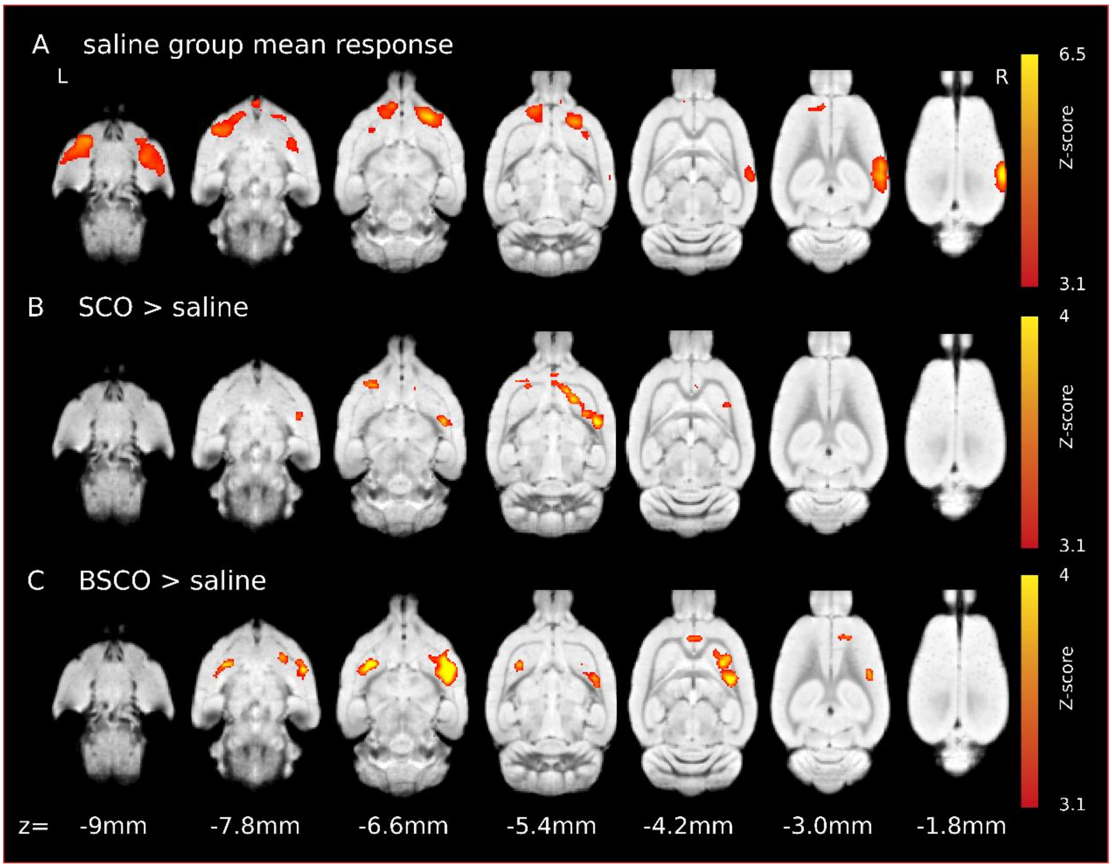


**Suppl. Fig. 12. The original investigation of the effect of SCO (1 mg/kg) and BSCO (1 mg/kg) by our group: left whisker stimulation evoked gradient-echo EPI sequence BOLD responses in isoflurane anesthesia.** We surprisingly observed increased BOLD activation in widespread frontal cortical areas. Mean activation evoked by whisker sensory stimulation in the barrel field ((distance from bregma): z = -9.0 mm, z = -7.8 mm, z = -6.6 mm, z = -5.4 mm, z = -4.2, z = -3.0, z = -2.6 mm). **A:** Sal; **B:** SCO > Sal and **C:** BSCO > Sal contrasts. Red-to-yellow colors represent suprathreshold Z scores, the corresponding colorbars depicted on the right.

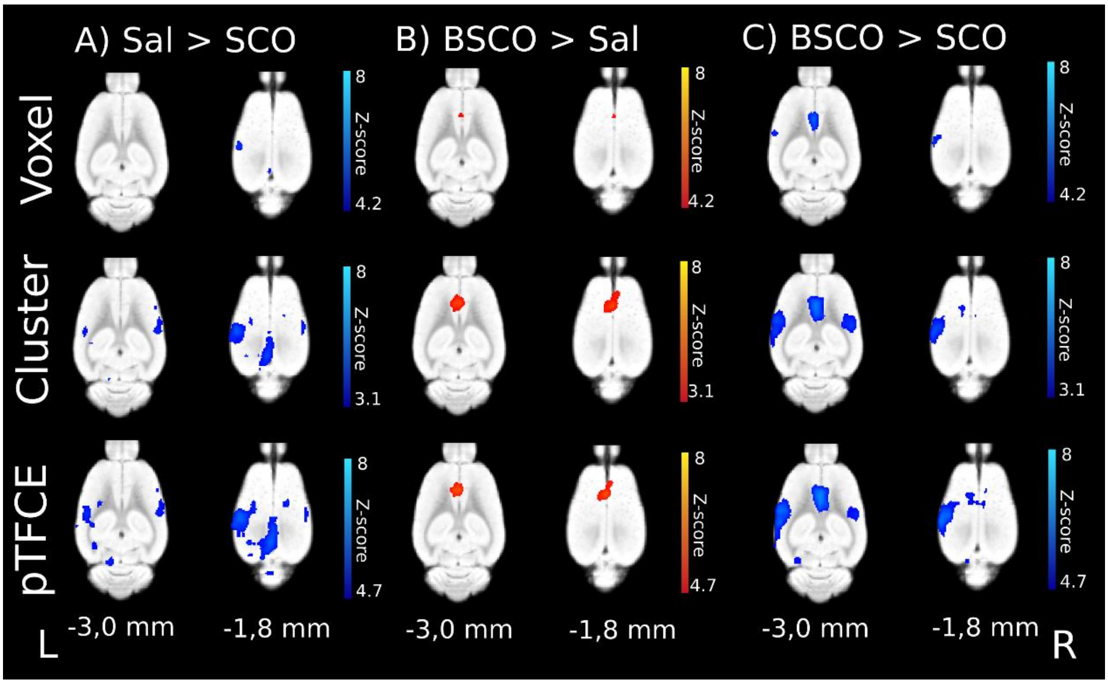


**Suppl. Fig. 13** Evoked BOLD response differences between treatment groups with GE EPI sequence in isoflurane anesthesia (n=18, 3*18). *(A)* Decreased BOLD response in the scopolamine (SCO) treated group compared to saline (Sal > SCO). *(B)* Increased BOLD response in the butylscopolamine (BSCO) group compared to saline (BSCO > Sal). *(c)* Differences between the SCO and BSCO treated groups (BSCO > SCO). The Z-scores in the upper row (Voxel) represent voxelwise statistical inference ($p<0.05$ correcting for multiple comparisons via GRF theory based FWER, navy blue-to-light blue or red-to-yellow from z=4.2 to z=8); in the middle row (Cluster) clusterwise statistical inference ($p<0.05$ correcting for multiple comparisons via GRF theory based FWER and a cluster forming threshold of p=0.001, navy blue-to-light blue or red-to-yellow z=3.1 to z=8); in the lower row probabilistic TFCE (pTFCE) statistics ($p<0.05$ correcting for multiple comparisons via GRF theory based FWER, navy blue-to-light blue or red-to-yellow z=4.7 to z=8) from the same data. The corresponding colorbars are depicted on the right. The z coordinates (distance from Bregma) of the selected slices (-3.0 mm; -1.8 mm) are given at the bottom of the figure (L, R stands for directions).

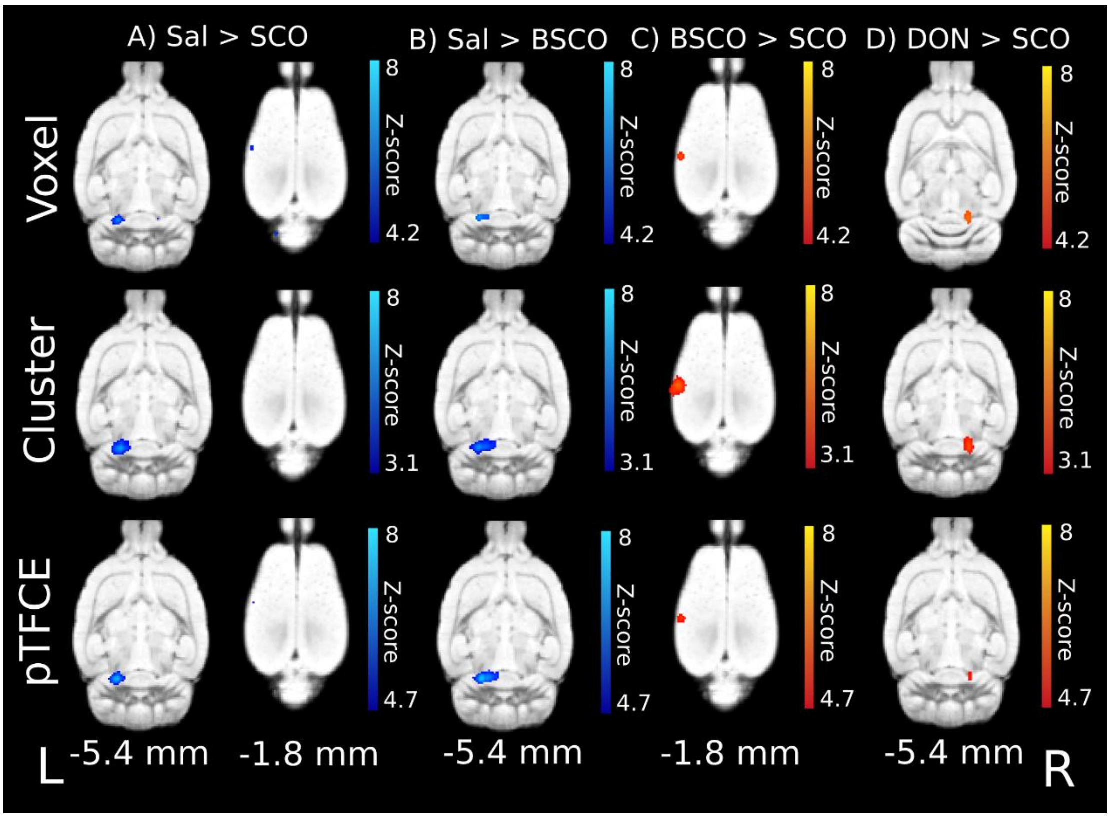


**Suppl. Fig. 14** Evoked BOLD response differences between groups with GE EPI sequence in medetomidine-isoflurane combined anesthesia (n=16, 2*15 – Sal, DON; 2*16 SCO, BSCO). *(A)* Sal > SCO contrast. *(B)* Sal > BSCO. *(C)* BSCO > SCO contrast. *(D)* DON > SCO contrast. Decreased BOLD responses in the BC due to SCO *(A, C)* and in the IC due to both SCO and BSCO *(A, B)*. Increased BOLD responses in the right IC in the DON treatment group compared to the SCO group *(D)*. The Z-scores in the upper row (Voxel) represent voxelwise statistical inference (p<0.05 correcting for multiple comparisons via GRF theory based FWER, navy blue-to-light blue or red-to-yellow from z=4.2 to z=8); in the middle row (Cluster) clusterwise statistical inference (p<0.05 correcting for multiple comparisons via GRF theory based FWER and a cluster forming threshold of p=0.001, navy blue-to-light blue or red-to-yellow z=3.1 to z=8); in the lower row probabilistic TFCE (pTFCE) statistics (p<0.05 correcting for multiple comparisons via GRF theory based FWER, navy blue-to-light blue or red-to-yellow z=4.7 to z=8) from the same data. The corresponding colorbars are depicted on the right. The z coordinates (distance from Bregma) of the selected slices (-5.4 mm; -1.8 mm) are given at the bottom of the figure (L, R stands for directions).

### Supplementary Tables

| *Exp. 1* | *Animals n =18* | *Weight at scan., (g)* | *Isoflurane dose (%)* | *Average motion (mm) GE* | *Average motion (mm) ASL* | *Respiration rate (1/min)* |
|---|---|---|---|---|---|---|
| **Sal** | *Mean (± sd)* | 335.94 (± 38.48) | 1.55 (± 0.11) | 0.004 (± 0.002) | 0.012 (± 0.003) | 72.48 (± 11.60) |
| | *Min* | 283 | 1.5 | 0.002 | 0.008 | 38 |
| | *Max* | 405 | 2 | 0.012 | 0.021 | 104 |
| **SCOP** | *Mean (± sd)* | 329.43 (± 34.77) | 1.53 (± 0.12) | 0.003 (± 0.001) | 0.011 (± 0.002) | 73.9 (± 11.32) |
| | *Min* | 276 | 1 | 0.002 | 0.007 | 41 |
| | *Max* | 397 | 2 | 0.006 | 0.0158 | 103 |
| **BSCOP** | *Mean (± sd)* | 344.17 (± 37.07) | 1.54 (± 0.13) | 0.004 (± 0.002) | 0.011 (± 0.003) | 73.32 (± 9.52) |
| | *Min* | 280 | 1.5 | 0.002 | 0.007 | 48 |
| | *Max* | 394 | 2 | 0.010 | 0.016 | 94 |

**Suppl. Table 1.** Summary statistics of the first experiment: weight, mean isoflurane dose, average in-scanner motion (root-mean-square (RMS) of relative displacement during GE and ASL imaging), and mean respiration rate of the rats (N=18).

| *Exp. 2* | *Animals n = 16* | *Weight at scan., (g)* | *Isoflurane dose (%)* | *Average motion (mm) GE* | *Average motion (mm) SE* | *Respiration rate (1/min)* |
|---|---|---|---|---|---|---|
| **Sal** | Mean (± sd) | 369.45 (± 47.94) | 0.77 (± 0.30) | 0.003 (± 0.001) | 0.006 (± 0.001) | 61.13 (±11.68) |
| | Min | 302 | 0.25 | 0.0018 | 0.005 | 42 |
| | Max | 454 | 1.2 | 0.0044 | 0.011 | 107 |
| **SCOP** | Mean (± sd) | 367.75 (± 38.95) | 0.74 (± 0.35) | 0.003 (± 0.002) | 0.007 (± 0.001) | 64.70 (± 13.43) |
| | Min | 288 | 0.1 | 0.002 | 0.005 | 40 |
| | Max | 433 | 1.5 | 0.008 | 0.009 | 100 |
| **BSCOP** | Mean (± sd) | 366.25 (± 28.86) | 0.68 (± 0.32) | 0.003 (± 0.001) | 0.006 (± 0.001) | 67.69 (± 12.85) |
| | Min | 324 | 0.25 | 0.002 | 0.005 | 47 |
| | Max | 416 | 1.5 | 0.007 | 0.010 | 100 |
| **DON** | Mean (± sd) | 364.89 (± 42.98) | 0.77 (± 0.33) | 0.006 (± 0.003) | 0.007 (± 0.002) | 59.53 (± 13.26) |
| | Min | 300 | 0.25 | 0.003 | 0.005 | 40 |
| | Max | 443 | 1.5 | 0.013 | 0.012 | 100 |

**Suppl. Table 2.** Summary statistics of the second experiment: in 0.005 mg/kg i.p. dexmedetomidine + isoflurane (0.5% - 1.5%). Weight, mean isoflurane dose, average in-scanner motion (mean RMS of relative displacement during GE and SE imaging), and mean respiration rate of the rats (N=16).

<table>
<tr><th rowspan="3">Study</th><th colspan="7">Voxelwise inference</th></tr>
<tr><th rowspan="2">Size of clusters (mm³)</th><th rowspan="2">Z-score of local maximums (3 peaks at most per cluster)</th><th rowspan="2">Signal change (%)</th><th colspan="3">Coordinates of local maximum</th><th rowspan="2">Structures to which local maximum Z-score belongs to:</th></tr>
<tr><th>x</th><th>y</th><th>z</th></tr>
<tr><td rowspan="3">Exp. 1, GE</td><td rowspan="3">622.6</td><td>13.8</td><td>0.24</td><td>-5.1</td><td>-3.2</td><td>-2.2</td><td>L S1 BF - Left primary somatosensory cortex, barrel field</td></tr>
<tr><td>10.8</td><td>0.16</td><td>0.3</td><td>4.8</td><td>-3.2</td><td>R PrL - Right prelimbic cortex</td></tr>
<tr><td>10.5</td><td>0.22</td><td>0.1</td><td>3.0</td><td>-3.4</td><td>R PrL - Right prelimbic cortex</td></tr>
<tr><td>Exp. 1, ASL</td><td>0.2</td><td>4.68</td><td>4.46</td><td>-4.9</td><td>-2.2</td><td>-2.4</td><td>L S1 BF - Left primary somatosensory cortex, barrel field</td></tr>
<tr><td rowspan="9">Exp. 2, GE</td><td rowspan="3">138.6</td><td>23.7</td><td>0.60</td><td>-2.3</td><td>-9.6</td><td>-4.8</td><td>L ECIC - Left external cortex of the inferior colliculus</td></tr>
<tr><td>15.8</td><td>0.17</td><td>1.7</td><td>-9.8</td><td>-4.4</td><td>R CIC - Right inferior colliculus</td></tr>
<tr><td>7.78</td><td>0.12</td><td>-3.3</td><td>-6.2</td><td>-6.2</td><td>L bic –Left brachium inferior colliculus</td></tr>
<tr><td rowspan="3">50.1</td><td>17.9</td><td>0.26</td><td>-5.1</td><td>-2.8</td><td>-2.2</td><td>L S1 BF - Left primary somatosensory cortex, barrel field</td></tr>
<tr><td>7.4</td><td>1.2</td><td>-7.1</td><td>-5.4</td><td>-4.4</td><td>L Au1 - Left primary auditory cortex</td></tr>
<tr><td>7.39</td><td>0.14</td><td>-6.9</td><td>-6</td><td>-5.6</td><td>L Au1 - Left primary auditory cortex</td></tr>
<tr><td rowspan="2">3.424</td><td>6.28</td><td>0.03</td><td>6.5</td><td>-5.6</td><td>-4.2</td><td>R Au1 - Right primary auditory cortex</td></tr>
<tr><td>5.3</td><td>0.12</td><td>6.9</td><td>-6.4</td><td>-5.4</td><td>R AuV - Right secondary auditory ventral cortex</td></tr>
<tr><td>0.1</td><td>4.56</td><td>0.46</td><td>2.5</td><td>-13</td><td>-8.2</td><td>R Sp5I – Spinal trigeminal nucleus, interpolar</td></tr>
<tr><td>Exp. 2, SE</td><td>4.4</td><td>7.09</td><td>0.39</td><td>-5.1</td><td>-2.8</td><td>-2</td><td>L S1 BF - Left primary somatosensory cortex, barrel field</td></tr>
</table>

**Suppl. Table 3.** The size and location of the fMRI BOLD response to whisker blow in the saline group in experiment 1. (GE, ASL) and experiment 2. (GE, SE). The size of each cluster, signal change (%) and the location (from Rat Brain Atlas of Paxinos and Watson) of Z-score (voxelwise statistical inference with $p<0.05$, correcting for multiple comparisons via voxel significance) local maximums (3 peaks at most per cluster) of BOLD response to somatosensory stimuli in saline groups.

| *Exp. 1, GE* | | | | | | | |
|---|---|---|---|---|---|---|---|
| *Treatment* | ***Voxelwise*** | | ***Clusterwise*** | | ***pTFCE*** | | |
| | *Size ($mm^3$)* | *Structures to which maximum Z-score belongs to:* | *Size ($mm^3$)* | *Structures to which maximum Z-score belongs to:* | *Size ($mm^3$)* | | *Structures to which maximum Z-score belongs to:* |
| | | | | | *Uncor-rected (3.1 < z)* | *Cor-rected (4.7 < z)* | |
| ***Sal*** | 622.6 | L S1 BF - Left primary somatosensory cortex, barrel field | 853.7 | L S1 BF - Left primary somatosensory cortex, barrel field | 1059.8 | 914.6 | L S1 BF - Left primary somatosensory cortex, barrel field |
| ***SCO*** | 413.5 | L PrL - Left prelimbic cortex | 624.4 | L PrL - Left prelimbic cortex | 781.0 | 667.7 | L PrL - Left prelimbic cortex |
| | 37.4 | R V1 B - Right primary visual cortex, binocular area | | | | | |
| ***BSCO*** | 605.8 | L S1 BF - Left primary somatosensory cortex, barrel field | 787.0 | L S1 BF - Left primary somatosensory cortex, barrel field | 1000.0 | 841.2 | L PrL - Left prelimbic cortex |

**Suppl. Table. 4.** The size of each cluster and the location (from Rat Brain Atlas of Paxinos and Watson) of Z-score maximums (Exp. 1, isoflurane anesthesia, GE) of the fMRI BOLD response to somatosensory stimuli in different treatment groups (saline, scopolamine, and butylscopolamine). Three statistical inferences (voxelwise with $p<0.05$, correcting for multiple comparisons via voxel significance; clusterwise corrected significance threshold of $p<0.05$ and cluster holding threshold of $3.1<Z$; uncorrected pTFCE-wise enhanced Z-scores thresholded at $p<0.001$, which corresponds to $3.1<Z$; corrected for multiple comparisons via voxel significance of $p<0.05$ of enhanced pTFCE Z-scores, which corresponds to $4.7<Z$) result in different cluster magnitude.

| *Exp. 1 GE* | | | | | | | |
|---|---|---|---|---|---|---|---|
| *Treatment* | *Size of voxelwise clusters ($mm^3$)* | *Z-score local maximum* | *Signal change (%):* | *Coordinates of local maximum* | | | *Structures to which z-score maximum peak belongs to:* |
| | | | | *x* | *y* | *z* | |
| ***Sal*** | 622.6 | 13.8 | 0.24 | -5.1 | -3.2 | -2.2 | L S1 BF - Left primary somatosensory cortex, barrel field |
| | | 10.8 | 0.16 | 0.3 | 4.8 | -3.2 | R PrL - Right prelimbic cortex |
| | | 9.32 | 0.10 | -6.3 | -3.6 | -6.6 | L Au1 - Left primary auditory cortex |
| | | 9.3 | -0.09 | -0.1 | 3.0 | -5.6 | L IL – Left infralimbic cortex |
| | | 9.12 | 0.44 | 6.5 | -7.0 | -5.2 | R TeA - Right temporal associatin cortex |
| | | 8.99 | -0.05 | 3.3 | 2.4 | -5.8 | R VO - Right ventral orbital cortex |
| ***SCO*** | 413.5 | 12.9 | 0.39 | -0.1 | 3.6 | -2.8 | L PrL - Left prelimbic cortex |
| | | 10.6 | 0.47 | -3.3 | 1 | -6.6 | L CPu – caudate putamen |
| | | 10.1 | 0.48 | 3.5 | 0.4 | -7.4 | R CPu - Right caudate putamen |
| | | 9.91 | 0.52 | 2.1 | 3.2 | -5.0 | R VO - Right ventral orbital cortex |
| | | 8.97 | 1.17 | -6.9 | -7.2 | -5.0 | L TeA -Left temporal associatin cortex |
| | | 8.84 | 0.45 | -5.3 | -3.4 | -2.2 | L S1 BF - Left primary somatosensory cortex, barrel field |
| | 37.4 | 7.18 | 0.79 | 5.9 | -7.4 | -3.6 | R V2L – Right secondary visual cortex |
| | | 5.59 | -0.05 | 2.5 | -9.8 | -2.8 | R prf cerebellum (near ecic) |
| | | 4.86 | 0.44 | 2.9 | -5.6 | -1.4 | R V2ML – Right secondary visual cortex, mediolateral |
| ***BSCO*** | 605.8 | 13.9 | 0.07 | -5.5 | -3.4 | -2.2 | L S1 BF - Left primary somatosensory cortex, barrel field |
| | | 12.4 | 0.03 | -0.7 | 3.6 | -4.0 | L PrL - Left prelimbic cortex |
| | | 10.8 | 0.4 | -2.7 | 0.4 | -6.8 | L CPu – Left caudate putamen |
| | | 10.7 | 0.09 | -6.5 | -3.2 | -6.4 | L AuV – Left secondary auditory ventral cortex |
| | | 10.6 | 0.22 | 2.3 | 2.6 | -5.4 | R VO – Right ventral orbital cortex |
| | | 10.3 | 0.26 | -6.9 | -5.6 | -4.4 | L Au1 - Left primary auditory cortex |

**Suppl. Table. 5.** The size of activations of fMRI BOLD and the location (from Rat Brain Atlas of Paxinos and Watson) of voxelwise Z-score local maximums (Exp. 1, GE). The size of activation clusters in different treatment groups (Sal, SCO, BSCO), signal change (%) in local maximums, and the location (from Rat Brain Atlas of Paxinos and Watson) of the six highest Z-score are shown. The minimum peak distance is set to 3 mm.

| *Exp. 1, GE* | | | | | | | |
|---|---|---|---|---|---|---|---|
| *Treat-ment* | *Voxelwise* | | *Clusterwise* | | *pTFCE* | | |
| | *Size ($mm^3$)* | *Structures to which maximum Z-score belongs to:* | *Size ($mm^3$)* | *Structures to which maximum Z-score belongs to:* | *Size ($mm^3$)* *Uncor-rected (3.1 < z)* | *Size ($mm^3$)* *Cor-rected (4.7 < z)* | *Structures to which maximum Z-score belongs to:* |
| **Sal > SCO** | 1.5 | L V1 M - Left primary visual cortex, monocular area | 20.6 | L V1 M - Left primary visual cortex, monocular area | 251.0 | 56.6 | L V1 M - Left primary visual cortex, monocular area |
| | 1.2 | L S1 BF - Left primary somatosensory cortex, barrel field | 14.3 | L S1 BF - Left primary somatosensory cortex, barrel field | 15.6 | - | R Ect - Right ectorhinal cortex |
| | 0.8 | L RSGc - Left retrosplenial granular cortex, c region | 8.0 | R S1 BF - Right primary somatosensory cortex, barrel field | 0.9 | - | R M1 - Right primary motor cortex |
| | 0.2 | R Ect - Right ectorhinal cortex | | | 0.1 | - | L VS - Left Ventral subiculum |
| | | | | | - | 8.7 | R S2 - Right secondary somatosensory cortex |
| | | | | | - | 4.1 | L PrS – Left presubiculum |
| | | | | | - | 0.4 | R M2 - RIght secondary motor cortex |
| **BSCO > Sal** | 1.5 | L Cg2 - Left cingulate cortex, area 2 | 9.3 | L Cg2 - Left cingulate cortex, area 2 | 35.8 | 7.6 | L Cg2 - Left cingulate cortex, area 2 |
| | | | | | 6.8 | - | L EpP - Left epipeduncular nucleus |
| | | | | | 0.4 | - | Left agranular insular cortex, ventral part |
| | | | | | 0.2 | - | L E - Left ependymal and subependymal |
| | | | | | 0.1 | - | Ventral pallidum |
| **BSCO > SCO** | 3.6 | L S1 BF - Left primary somatosensory cortex, barrel field | 16.5 | L S1 BF - Left primary somatosensory cortex, barrel field | 288.3 | 28.7 | L S1 BF - Left primary somatosensory cortex, barrel field |
| | 2.5 | L Cg2 - Left cingulate cortex, area 2 | 12.5 | L Cg2 - Left cingulate cortex, area 2 | 0.1 | - | R V1 B - Right primary visual cortex, binocular area |
| | | | 7.0 | R S1 BF - Right primary somatosensory cortex, barrel field | - | 15.8 | L Cg2 - Left cingulate cortex, area 2 |
| | | | | | - | 4.0 | L CPu – Left caudate putamen |
| | | | | | - | 3.3 | R S1 BF - Right primary somatosensory cortex, barrel field |
| | | | | | - | 1.3 | L PRh – Left perirhinal cortex |
| | | | | | - | 0.6 | R V1 B - Right primary visual cortex, monocular area |
| **Sal > BSCO** | 0.5 | R FrA - Right frontal association cortex | | | 9.2 | 0.5 | R FrA - Right frontal assocn cortex |
| | 0.2 | R FrA - Right frontal association cortex | | | 7.4 | 0.1 | L V1 M -Left primary visual cortex, monocular area |
| | 0.1 | L FrA -Left frontal association cortex | | | 3 | - | R PRh - Right perirhinal cortex |
| | | | | | 2.9 | - | L M2 - Left secondary motor cortex |
| | | | | | 2.5 | - | R M2 - Right secondary motor cortex |
| | | | | | 2.4 | - | R M1 - Right primary motor cortex |
| | | | | | 2.9 | - | R LO - Right lateral orbital cortex |
| | | | | | 0.7 | - | L Cl -Left Cerebellar lobes |
| | | | | | 0.2 | - | L RSD - Left retrosplenial dysgranular cortex |
| | | | | | - | 0.1 | R M2 - Right secondary motor cortex |
| **SCO > Sal** | | | | | 0.2 | - | L Cg1 - Left cingulate cortex, area 1 |
| | 0.4 | R DLO - Right dorsolateral orbital cortex | | | 0.8 | - | R DLO - Right dorsolateral orbital cortex |

| | | | | | | |
|---|---|---|---|---|---|---|
| **SCO> BSCO** | 0.1 | L M2 – Left secondary motor cortex | | 0.3 | - | L M2 – Left secondary motor cortex |
| | 0.1 | L M2 - Left secondary motor cortex | | 0.2 | - | L M2 - Left secondary motor cortex |

**Suppl. Table 6.** The size of significant evoked BOLD activation differences between treatment groups with gradient-echo (GE) EPI sequence in isoflurane anesthesia (Exp. 1, GE). Different statistical inferences result in different cluster sizes and different cerebral structures of maximum peaks. The anatomical areas were identified using the standard orientation of the digitalized version of the Rat Brain Atlas of Paxinos and Watson.

| *Exp. 1, ASL* | | | | | | | | |
|---|---|---|---|---|---|---|---|---|
| *Treatment* | *Size of pTFCE-wise clusters, uncorrected Z > 3.1 (mm³)* | *Size of pTFCE-wise clusters, corrected Z > 4.7 (mm³)* | *Z-score of local maximum* | *Signal change (%)* | *Coordinates of local maximum* | | | *Structures to which local Z-score maximum belongs to:* |
| | | | | | *x* | *y* | *z* | |
| ***Sal*** | 6.0 | 0.1 | 4.78 | 4.45 | -4.9 | -2.2 | -2.4 | L S1 BF - Left primary somatosensory cortex, barrel field |
| | 1.2 | - | 3.62 | 6.40 | -2.1 | -8.4 | -4.2 | L ECIC - Left external cortex of the inferior colliculus |
| ***SCO*** | 3.0 | - | 3.88 | 4.23 | -4.5 | -3.0 | -2.4 | L S1 BF - Left primary somatosensory cortex, barrel field |
| | 1.0 | - | 3.87 | 7.05 | -2.9 | -7.2 | -4.6 | L bic - Left brachium of the inferior colliculus |
| | 0.1 | - | 3.35 | 1.12 | 0.3 | -6.4 | -1.8 | R RSGb - Left retrosplenial granular cortex, b region |
| ***BSCO*** | 87.2 | 0.2 | 4.86 | 4.27 | -4.9 | -2.8 | -2.0 | L S1 BF - Left primary somatosensory cortex, barrel field |
| | | - | 4 | 10.67 | -7.1 | -5.2 | -5 | L Au1 - Left primary auditory cortex |
| | | - | 3.98 | 2.56 | 5.7 | -5.6 | -3.4 | L V2L - Left secondary visual cortex |
| | | - | 3.96 | 4.95 | -1.7 | 0.6 | -1.4 | L M1 - Left primary motor cortex |
| | | - | 3.94 | 3.20 | 1.7 | 1.8 | -1.6 | R M1 - Right primary motor cortex |
| | | - | 3.82 | 2.76 | 4.1 | -0.8 | -1.6 | R S1 BF - Right primary somatosensory cortex, barrel field |
| | 7.3 | - | 4.03 | 3.30 | -2.5 | -8.0 | -4.4 | L ECIC - Left external cortex of the inferior colliculus |
| | 2.8 | - | 4.25 | 1.92 | 1.7 | -7.6 | -4.6 | R DpWh –Right deep white layer superior colliculus |
| | 0.5 | - | 3.38 | 5.72 | -2.3 | -7.8 | -9.2 | L VLL - Ventral nucleus of lateral lemniscus |

**Suppl. Table 7.** The size and location of the local Z-score maximums of fMRI BOLD response to whisker blow in ASL. The size of each cluster, signal change (%) and the location (from Rat Brain Atlas of Paxinos and Watson) of local maximums (pTFCE-wise enhanced uncorrected

[3.1<z] and GRF corrected [p<0.05, which corresponds to 4.7<z] Z-scores) of BOLD response to somatosensory stimuli in different treatment groups. The minimum peak distance is set to 3.0 mm.

| *Exp. 2, GE* | | | | | | | |
|---|---|---|---|---|---|---|---|
| *Treatment* | *Size of voxelwise clusters (mm³)* | *Z-score local maximum* | *Signal change (%):* | *Coordinates of local maximum* | | | *Structures to which z-score maximum peak belongs to:* |
| | | | | *x* | *y* | *z* | |
| **Sal** | 138.6 | 23.70 | 0.60 | -2.3 | -9.6 | -4.8 | L ECIC - Left external cortex of the inferior colliculus |
| | | 15.80 | 0.17 | 1.7 | -9.8 | -4.4 | R CIC - Right central nu inferior colliculus |
| | | 7.78 | 0.18 | -3.3 | -6.2 | -6.2 | L bic –Left brachium inferior colliculus |
| | | 7.04 | 0.28 | 3.3 | -7.4 | -6.2 | R bic – Right brachium inferior colliculus |
| | | 5.39 | 0.29 | -3.3 | -4.4 | -6.4 | L str – Left superior thalamic radiation |
| | | 4.82 | 0.35 | -4.1 | -10.2 | -2.8 | Left cerebellar lobes |
| | 50.1 | 17.90 | 0.26 | -5.1 | -2.8 | -2.2 | L S1 BF - Left primary somatosensory cortex, barrel field |
| | | 7.40 | 1.21 | -7.1 | -5.4 | -4.4 | L Au1 - Left primary auditory cortex |
| | | 7.39 | 0.14 | -6.9 | -6 | -5.6 | L Au1 - Left primary auditory cortex |
| | | 7.31 | 1.22 | -7.1 | -4.4 | -4.4 | L Au1 - Left primary auditory cortex |
| | 3.4 | 6.28 | 0.11 | 6.5 | -5.6 | -4.2 | R Au1 – Right primary auditory cortex |
| | | 5.30 | 0.12 | 6.9 | -6.4 | -5.4 | R AuV – Right secondary auditory cortex, ventral |
| | 0.1 | 4.56 | 0.46 | 2.5 | -13 | -8.2 | R Sp5I – Spinal trigeminal nucleus, interpolar |
| **SCO** | 140.6 | 19.30 | 0.20 | -2.3 | -9.4 | -5 | L ECIC - Left external cortex of the inferior colliculus |
| | | 18.60 | 0.06 | -1.9 | -9.8 | -4.2 | L CIC - Left central nu inferior colliculus |
| | | 12.10 | -0.45 | 1.9 | -10.2 | -4 | Right cerebellum |
| | | 7.12 | 0.23 | -3.1 | -7.2 | -6.2 | L str – Left superior thalamic radiaion |
| | | 6.42 | 0.07 | 3.7 | -6.6 | -6.6 | R PrS – Right presubiculum |
| | 39.5 | 13.50 | 0.20 | -5.1 | -2.6 | -2.4 | L S1 BF - Left primary somatosensory cortex, barrel field |
| | | 7.38 | 0.56 | -6.7 | -5.8 | -4.4 | L Au1 - Left primary auditory cortex |
| | | 7.38 | 0.48 | -7.1 | -5.2 | -5.8 | L TeA - Left temporal associatin cortex |
| | | 6.00 | 0.78 | -6.7 | -3.4 | -4.4 | L Au1 - Left secondary auditory cortex, dorsal |
| | 2.1 | 5.41 | 0.44 | 6.7 | -6.6 | -5.2 | R TeA - Right temporal associatin cortex |
| | | 5.40 | 0.51 | 6.5 | -6.2 | -4.8 | R Au1 - Right secondary auditory cortex, dorsal |
| | | 5.39 | 0.73 | 6.7 | -5.6 | -4.6 | R Au1 - Right primary auditory cortex |
| | 0.4 | 4.76 | 0.29 | -2.9 | -4.6 | -7.2 | L ZIC -Left zona incerta, caudal part |
| | 0.3 | 4.94 | -0.87 | 5.1 | -12.2 | -4.4 | Right celebellar lobes |
| **BSCO** | 126.8 | 21.1 | 0.14 | -2.3 | -9.6 | -4.8 | L ECIC - Left external cortex of the inferior colliculus |
| | | 12.9 | -0.32 | 1.9 | -9.8 | -4.2 | R ECIC - Right external cortex of the inferior colliculus |
| | | 6.77 | -0.04 | -3.5 | -6.4 | -6 | L MGV - |
| | 49.8 | 19.2 | 0.51 | -5.3 | -3 | -2 | L S1 BF - Left primary somatosensory cortex, barrel field |
| | | 6.39 | 0.89 | -7.3 | -4.6 | -6 | L TeA - Left temporal associatin cortex |

| | | | | | | | |
|---|---|---|---|---|---|---|---|
| | 0.8 | 5.11 | 0.33 | 5.3 | -2.6 | -2.8 | R S1 BF - Right primary somatosensory cortex, barrel field |
| ***DON*** | 95.6 | 19.2 | 0.21 | -2.1 | -9.6 | -4.6 | L ECIC - Left external cortex of the inferior colliculus |
| | | 16 | 0.16 | 1.9 | -9.8 | -4.6 | R ECIC - Right external cortex of the inferior colliculus |
| | | 5.77 | 0.34 | -3.5 | -6.8 | -6 | L MG – Left medial geniculate nucleus |
| | | 4.76 | -0.23 | 2.5 | -12.2 | -7.6 | Right cerebellar lobes |
| | 29.4 | 14 | 0.06 | -5.1 | -2.4 | -2.2 | L S1 BF - Left primary somatosensory cortex, barrel field |
| | | 6.19 | 0.02 | -6.7 | -5.2 | -4.4 | L Au1 - Left primary auditory cortex |
| | | 6.17 | 0.00 | -6.5 | -4.6 | -4.2 | L Au1 - Left primary auditory cortex |
| | | 5.46 | 0.19 | -7.1 | -5.2 | -5.8 | L TeA - Left temporal associatin cortex |

**Suppl. Table 8.** The size and location of the local Z-score maximums of fMRI BOLD response to whisker blow in combined anesthesia (GE). The size of each cluster, signal change (%), and the location (from Rat Brain Atlas of Paxinos and Watson) of voxelwise Z-score (correcting for multiple comparisons via voxel significance of $p<0.05$) local maximums of BOLD response to somatosensory stimuli in different treatment groups. The minimum peak distance is set to 0.5 mm.

| *Exp. 2, SE* | | | |
|---|---|---|---|
| *Treatment* | *Size of voxelwise clusters (mm³)* | *Structures to which voxelwise Z-score peak belongs to:* | *Average signal increase in the related structures (%):* |
| ***Sal*** | 4.4 | L S1 BF - Left primary somatosensory cortex, barrel field | 0.38 |
| ***SCO*** | 1.3 | L S1 BF - Left primary somatosensory cortex, barrel field | 0.50 |
| ***BSCO*** | 6.3 | L S1 BF - Left primary somatosensory cortex, barrel field | 0.15 |
| ***DON*** | 8.4 | L S1 BF - Left primary somatosensory cortex, barrel field | 0.02 |

**Suppl. Table 9.** The size ($mm^3$) and location of the evoked mean BOLD response in saline, scopolamine, butylscopolamine, and donepezil groups with SE sequence in isoflurane-medetomidine combined anesthesia. Rat Brain Atlas of Paxinos and Watson define structures to which voxelwise Z-score (correcting for multiple comparisons via voxel significance of $p<0.05$) peaks belong to. Signal change in the structure is defined as the ratio of the average signal during the whisker stimulation and resting periods.

| *Exp. 2, GE* | | | | | | |
|---|---|---|---|---|---|---|
| | *Voxelwise* | | *Clusterwise* | | *pTFCE* | |
| *Treatment* | *Size of clusters (mm$^3$)* | *Structures to which maximum Z-score belongs to:* | *Size of clusters (mm$^3$)* | *Structures to which maximum Z-score belongs to:* | *Size of clusters (mm$^3$)* | *Structures to which maximum Z-score belongs to:* |
| **Sal > SCO** | 3.9 | L ECIC- Left external cx inferior colliculus | 10.6 | L ECIC- Left external cx inferior colliculus | 14.2 | L ECIC- Left external cx inferior colliculus |
| | 0.1 | L S1 BF- Left primary somatosensory cortex, barrel field | | | 5.1 | R ECIC- Right external cx inferior colliculus |
| | | | | | 4.0 | L S1 BF- Left primary somatosensory cortex, barrel field |
| | | | | | 0.1 | L cerebellar lobes (p.a.) |
| **Sal > BSCO** | 4.1 | L ECIC- Left external cx inferior colliculus | 15.2 | L cerebellar lobes (expanded from L ECIC) | 33.6 | L ECIC- Left external cx inferior colliculus |
| | 1.3 | L cerebellar lobes (p.a.) | | | | |
| **Sal > DON** | | | 4.0 | L cerebellar lobes (expanded from L ECIC) | 20.8 | L cerebellar lobes (expanded from L ECIC) |
| **SCO > Sal** | 0.3 | L cerebellar lobes (p.a.) | | | 4.0 | L cerebellar lobes (p.a.) |
| | | | | | 0.3 | R cerebellar lobes (p.a.) |
| | | | | | 0.2 | R cerebellar lobes (p.a.) |
| **SCO > BSCO** | | | | | 0.8 | L cerebellar lobes (p.a.) |
| **SCO > DON** | | | | | 3.1 | R cerebellar lobes (p.a.) - Crus2 |
| | | | | | 0.2 | L Cerebellar lobes (p.a.) |
| | | | | | 0.1 | L Vtg – Left ventral tegmental nucleus |
| **BSCO > Sal** | | | | | 5.2 | L Cerebellar lobes |
| **BSCO > SCO** | 0.4 | L S1 BF- Left primary somatosensory cortex, barrel field | 4.1 | L S1 BF- Left primary somatosensory cortex, barrel field | 5.9 | L S1 BF- Left primary somatosensory cortex, barrel field |
| **BSCO > DON** | | | | | | |
| **DON > Sal** | | | | | | |
| **DON > SCO** | 1.2 | R ECIC- Right external cx inferior colliculus | 5.5 | R ECIC- Right external cx inferior colliculus | 6.8 | R ECIC- Right external cx inferior colliculus |
| **DON > BSCO** | 1.0 | R cerebellar lobes (expanded from R ECIC) | 4.8 | R cerebellar lobes (expanded from R ECIC) | 7.0 | R cerebellar lobes (expanded from R ECIC) |
| | | | | | 2.1 | R cerebellar lobes (p.a.) |

**Suppl. Table 10.** Evoked activation differences (the size of clusters in mm$^3$ and the structures to which Z-score peaks belong) of fMRI BOLD response to whisker blow with GE sequence between treatment groups by whisker stimuli. Rat Brain Atlas of Paxinos and Watson define the structures. Between parentheses, the abbreviation p. a. stands for presumable artifact: our gradient-echo images are not accurate enough in the cerebellum to conclude anything from these results.

Andersson, J. L. R., Jenkinson, M., & Smith, S. (2007). *Non-linear optimisation.* Retrieved from Oxford:

Behzadi, Y., Restom, K., Liau, J., & Liu, T. T. (2007). A component based noise correction method (CompCor) for BOLD and perfusion based fMRI. *Neuroimage, 37*(1), 90-101. doi:10.1016/j.neuroimage.2007.04.042

Eklund, A., Nichols, T. E., & Knutsson, H. (2016). Cluster failure: Why fMRI inferences for spatial extent have inflated false-positive rates. *Proc Natl Acad Sci U S A, 113*(28), 7900-7905. doi:10.1073/pnas.1602413113

Friston, K. J., Worsley, K. J., Frackowiak, R. S., Mazziotta, J. C., & Evans, A. C. (1994). Assessing the significance of focal activations using their spatial extent. *Hum Brain Mapp, 1*(3), 210-220. doi:10.1002/hbm.460010306

Gao, J. H., & Liu, H. L. (2012). Inflow effects on functional MRI. *Neuroimage, 62*(2), 1035-1039. doi:10.1016/j.neuroimage.2011.09.088

Jenkinson, M., Bannister, P., Brady, M., & Smith, S. (2002). Improved optimization for the robust and accurate linear registration and motion correction of brain images. *Neuroimage, 17*(2), 825-841.

Jenkinson, M., Beckmann, C. F., Behrens, T. E., Woolrich, M. W., & Smith, S. M. (2012). Fsl. *Neuroimage, 62*(2), 782-790. doi:10.1016/j.neuroimage.2011.09.015

Jenkinson, M., & Smith, S. (2001). A global optimisation method for robust affine registration of brain images. *Med Image Anal, 5*(2), 143-156.

Nasrallah, F. A., Lee, E. L., & Chuang, K. H. (2012). Optimization of flow-sensitive alternating inversion recovery (FAIR) for perfusion functional MRI of rodent brain. *NMR Biomed, 25*(11), 1209-1216. doi:10.1002/nbm.2790

Paxinos, G., & Watson, C. (Eds.). (2005). *The Rat Brain in Stereotaxic Coordinates*. Burlington, MA: Elsevier Academic Press.

Smith, S. M. (2002). Fast robust automated brain extraction. *Hum Brain Mapp, 17*(3), 143-155. doi:10.1002/hbm.10062

Smith, S. M., & Brady, J. M. (1997). SUSAN—A New Approach to Low Level Image Processing. *International Journal of Computer Vision, 23*(1), 45-78. doi:10.1023/a:1007963824710

Smith, S. M., & Nichols, T. E. (2009). Threshold-free cluster enhancement: addressing problems of smoothing, threshold dependence and localisation in cluster inference. *Neuroimage, 44*(1), 83-98. doi:10.1016/j.neuroimage.2008.03.061

Spisak, T., Pozsgay, Z., Aranyi, C., David, S., Kocsis, P., Nyitrai, G., . . . Kincses, Z. T. (2017). Central sensitization-related changes of effective and functional connectivity in the rat inflammatory trigeminal pain model. *Neuroscience, 344*, 133-147. doi:10.1016/j.neuroscience.2016.12.018

Spisak, T., Spisak, Z., Zunhammer, M., Bingel, U., Smith, S., Nichols, T., & Kincses, T. (2018). Probabilistic TFCE: A generalized combination of cluster size and voxel intensity to increase statistical power. *Neuroimage, 185*, 12-26. doi:10.1016/j.neuroimage.2018.09.078

van der Zwaag, W., Marques, J. P., Lei, H., Just, N., Kober, T., & Gruetter, R. (2009). Minimization of Nyquist ghosting for echo-planar imaging at ultra-high fields based on a "negative readout gradient" strategy. *J Magn Reson Imaging, 30*(5), 1171-1178. doi:10.1002/jmri.21951

Woolrich, M. W., Ripley, B. D., Brady, M., & Smith, S. M. (2001). Temporal autocorrelation in univariate linear modeling of FMRI data. *Neuroimage, 14*(6), 1370-1386. doi:10.1006/nimg.2001.0931

Worsley, K. J. (2001). Statistical analysis of activation images. In P. Jezzard, P. M. Matthews, & S. M. Smith (Eds.), *Functional MRI: An Introduction to Methods* (pp. 251-271). Oxford: Oxford Universtiy Press.
